# Applying functional MRI analysis techniques to whole-brain calcium-imaging data to investigate structural and functional relationships in *Drosophila melanogaster*

**DOI:** 10.1101/2025.07.01.662601

**Authors:** Takuto Okuno, Alexander Woodward, Hideyuki Okano, Junichi Hata

## Abstract

Clarifying the relationship between structure and function is important for understanding the brain. In *Drosophila melanogaster*, FlyEM and FlyWire electron microscopy-based connectome data and whole-brain calcium imaging data are available. We applied pre-processing methods from fMRI to whole-brain calcium imaging data and comprehensively investigated the optimal parameters. Then, we found that the FC-SC (functional and structural connectivity) correlation decreased linearly with region of interest count, and this trend was the same in flies and humans. We also developed a new, more robust method to quantify the degree of pre-and post-synaptic segregation and investigated this in the fly whole-brain. This revealed that many neurons have unsegregated synapses. We extracted highly unsegregated synapses and compared them with random-extracted null SC matrices. Their FC-SC correlation was significantly higher, indicating that these synapses contribute to FC well. Conversely, highly segregated-synapses showed significantly lower FC-SC correlation and contribute less to FC. Neurons with unsegregated synapses like non-spiking neurons are spread throughout the whole-brain, and they are thought to have a significant influence on FC.

## Introduction

Functional connectivity (FC) describes the relationship between functional activity in one brain region and that in another. In previous approaches, two spike trains were analyzed by a cross-correlation histogram [1, 2]; then, this method was applied to positron emission tomography and functional magnetic resonance imaging (fMRI) [3]. Subsequently, the time-series correlation between two regions has become widely used as a measure of undirected FC [4]. Underlying such functional connectivity, a structural connectivity (SC) substrate exists, and its relationship with FC has been investigated using various modalities [5, 6, 7, 8, 9, 10, 11]. In humans, FC-SC correlations between resting-state (rs-) fMRI and diffusion tensor imaging have been explored, and studies have shown high relationships of r=0.66 for 66 regions and r=0.54 for 998 regions [6, 7]. Other studies have examined the relationship between rs-fMRI and SC based on retrograde tracing from 145 injection sites in the left cortex of marmosets (r=0.379 for 9862 voxels, Ext.Data.Fig.2-1c) [10] and the relationship between calcium imaging and SC based on FlyEM connectome data of *Drosophila melanogaster (D. melanogaster)* [12] (r=0.74 for 37 regions) [9]. Because of the disparity in species, region of interest (ROI) count, and modality, a direct comparison across them becomes difficult. Therefore, in this study, we used electron microscopy (EM)-based FlyEM [12] and FlyWire [13] connectome data, which are currently considered to be the most detailed, and whole-brain calcium imaging data [14] of *D*. *melanogaster* to comprehensively investigate the relationship between FC and SC across various ROI shapes and counts.

High-quality whole-brain calcium imaging data have recently been published in [14] and, to our knowledge, is the only available open dataset. Therefore, an exact registration process between this type of functional data and compatible structural data has not been sufficiently investigated. Therefore, we applied registration procedures used in fMRI, quantified by FC-SC correlation and FC-SC detection, and investigated their optimality. Specifically, we investigated the application of motion correction and slice timing correction, as used in SPM [15], the application of nuisance factor removal methods [10], and the application of spatial smoothing to improve the accuracy of 2nd-level (group) analysis [16], aiming to calculate FC more accurately. This analysis revealed a relationship between ROI count and FC-SC correlation in the fly brain, and we further explored interspecies differences in FC-SC correlation between humans and marmosets.

Because of the strong relationship between FC and SC, they can be used to investigate inter-individual variability in *D*. *melanogaster*. FC varied across the eight flies in our dataset, and spatial smoothing can improve group analysis. Furthermore, SC also varied between FlyWire and FlyEM connectome data, and existing comparative studies [17, 18] have observed large differences in cell type classification [18]. Since these two EM connectomes have not been quantitatively compared to FC, we compared them based on the averaged FC of eight flies. FlyEM connectome data has a “confidence” threshold that indicates the certainty of an annotated synapse. Conversely, FlyWire has a “CleftScore” [19] threshold that shows the degree of synaptic detection. These initial thresholds adopted by FlyWire and FlyEM resulted in substantial differences in post-synaptic counts. We comprehensively investigated the impact of these thresholds by using FC-SC correlation and FC-SC detection in the EM connectome data of each fly.

In calcium imaging, the pre-synaptic activity in each region is detected as a calcium signal [20] representing functional activity. In textbook terms, post-synaptic sites on dendrites sense neurotransmitters and graded potentials if they are excitatory; and pre-synapses on axons release neurotransmitters [21, 22]. Then the FC is detected as the synchronization of functional activities. However, the synaptic arrangement on the neurons revealed in the EM connectome shows pre-and post-synapses that are highly intermingled in many neurons (i.e. input and output locations may be intermingled across axon and dendritic arbors); thus, signal transmission may be complex [22, 17, 13, 23]. Nerve cells include non-spiking neurons such as visual neurons, CT1 amacrine cells [24, 25, 26, 27, 28], interneurons in the antennal lobe [29], and APL cells in the mushroom body [30, 31, 32]. In *D*. *melanogaster*. these cells show localized functional activity taking place on a portion of the dendrite. In CT1 and APL cells, the pre-and post-synapses are highly intermingled (Fig.4a, Ext.Data.Fig.4-2). Therefore, we wanted to investigate whether these unsegregated synaptic neurons exist in the brain and we defined the pre-and post-synapse segregation index (PPSSI) to measure the degree of synaptic segregation for neurons in the whole brain. We also investigated the effect of such unsegregated synapses on FC. Unsegregated synapses are found not only in flies but also in mammals [33, 34, 21, 35, 36, 37], including humans (Ext.Data.Fig.1-2 from [38]). Of note, retinal amacrine cells have extensive reciprocal synapses [33, 34, 39, 40, 41], which provide local feedback inhibition of burst inputs [34]. This microscopic function in the retina, specifically compartmentalization via unsegregated synapses, may be generally applicable to the entire brain. The relationship between unsegregated synapses and functional connectivity could support this idea.

**Fig. 1.**
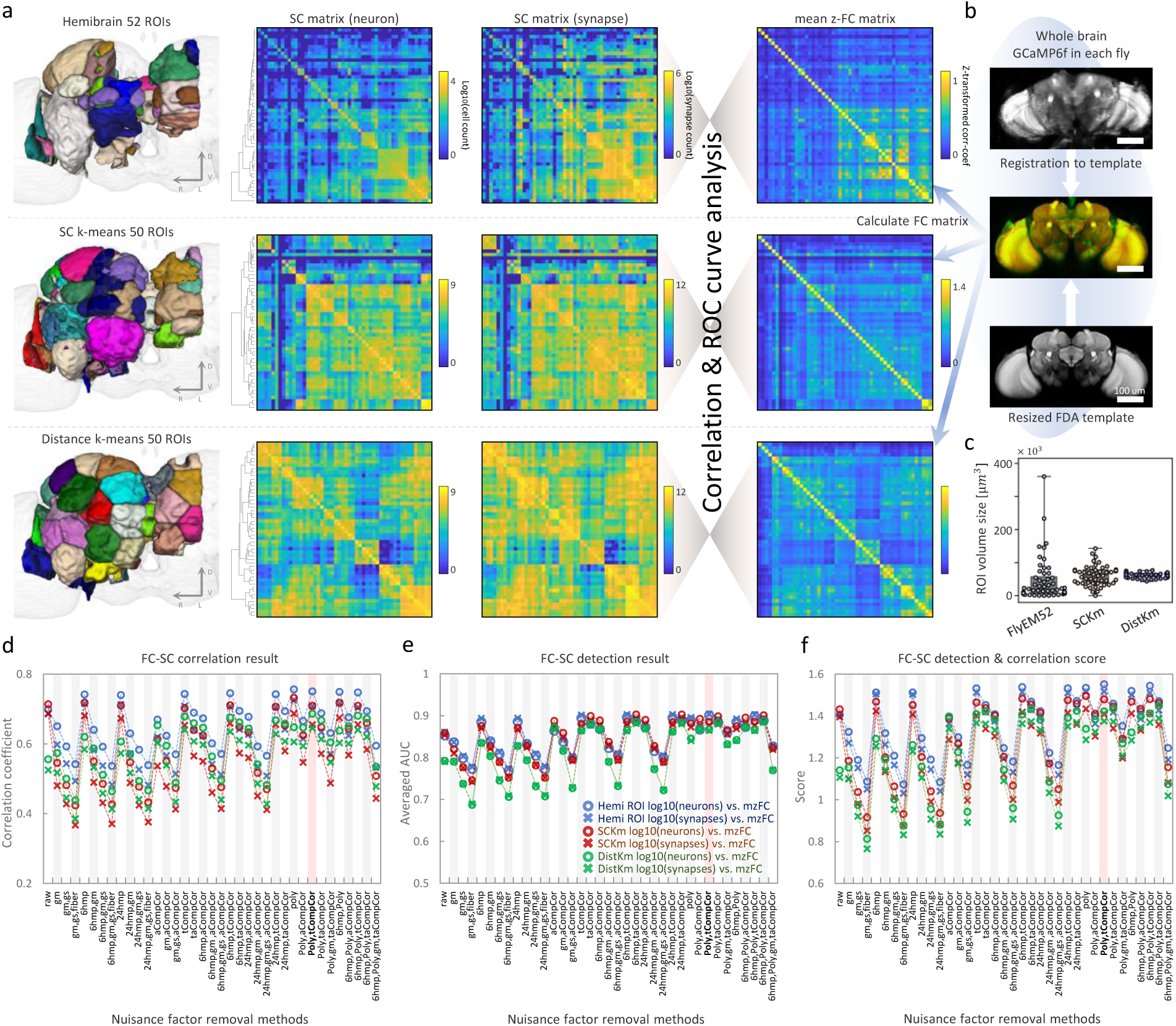
Comparison between SC and FC, and optimal nuisance factor removal methods. a, Schematic image of the comparison between SC matrices and FC matrices. (left) Three ROI types: Hemibrain anatomical 52 ROIs, SCKm 50 ROIs, and DistKm 50 ROIs. (center) SC matrices (neuron) and SC matrices (synapse) for each ROI type. (right) FC matrices for an average of 8 flies for each ROI type. **b,** Schematic image of calcium imaging data registration. **c**, Swarm plot of ROI volume for each ROI type. **d**, Results of nuisance factor removal method investigation. The vertical axis shows the correlation coefficient, and the horizontal axis shows the FC-SC correlation for each method. Results are plotted for the three ROI types. **e,** The vertical axis shows the averaged AUC of 100 thresholds, and the horizontal axis is the same as **d. f**, The vertical axis shows the FC-SC Detection & Correlation score, and the horizontal axis is the same as **d.**

**Fig. 2.**
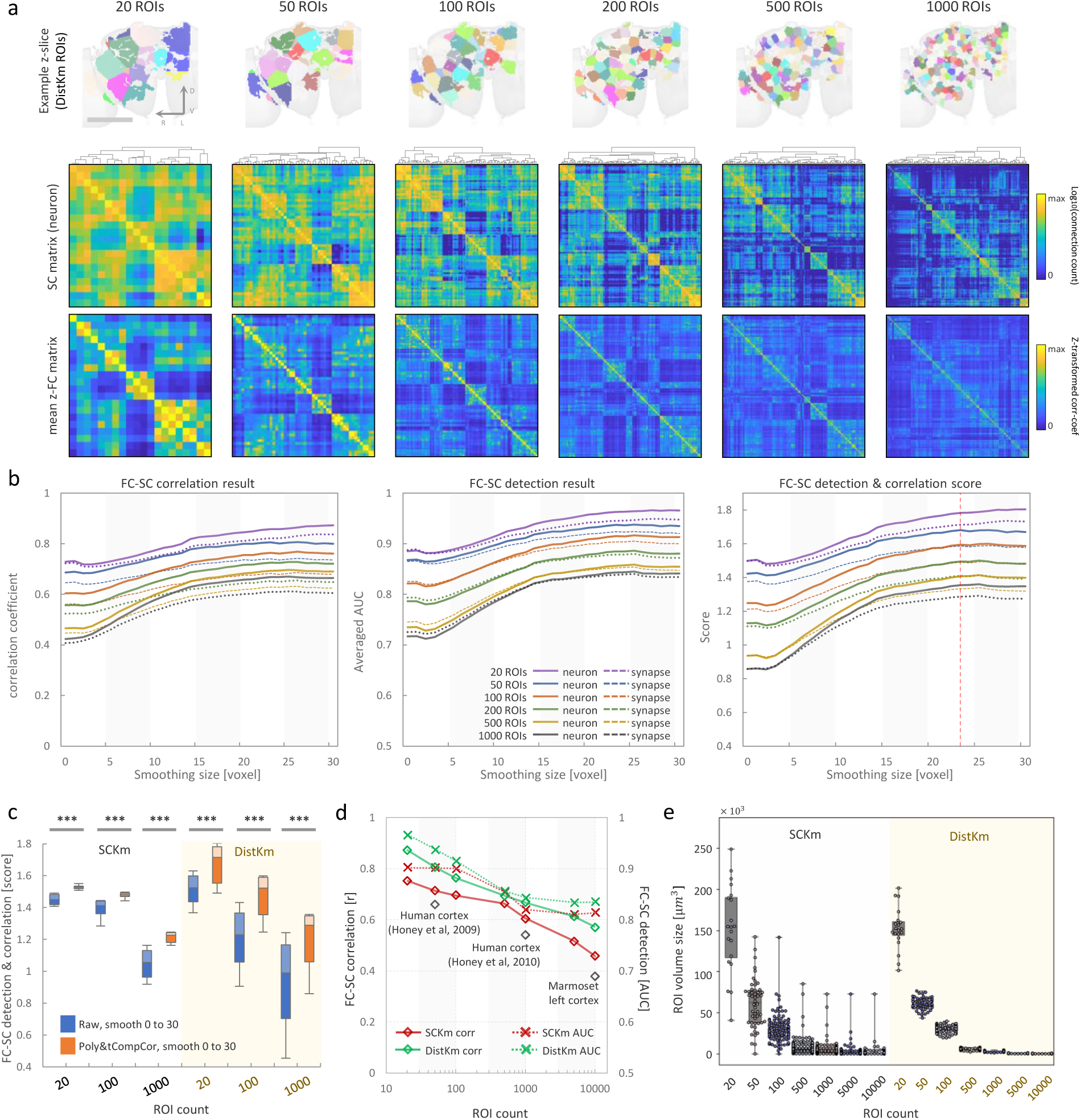
Investigating spatial smoothing size, and comparison of FC-SC correlation between mammals and flies. a, (top) Example z-slice of the reduced FDA template with DistKm ROIs from 20 to 1000 ROIs. (middle) SC matrices (neuron) of DistKm ROIs from 20 to 1000 ROIs. (bottom) FC matrices of DistKm ROIs from 20 to 1000 ROIs. **b,** From left to right, FC-SC correlation, FC-SC detection and FC-SC Detection & Correlation score, respectively. The solid line shows the FC vs. SC matrix (neuron), and the dashed line shows FC vs. SC matrix (synapse) at each ROI count. The vertical axis shows the correlation coefficient, averaged AUC and FC-SC Detection & Correlation score, respectively. The horizontal axis shows spatial smoothing size (voxels). **c,** Comparison between raw & smoothing (0 to 30 voxels) vs. Poly-tCompCor & smoothing (0 to 30 voxels) at each ROI count. Non-parametric Mann-Whitney U test was performed and Bonferroni correction was applied to correct for the familywise error rate (*** p<1.67e-4). **d,** Relation between FC-SC correlation and ROI count. (left vertical axis) Solid line shows FC-SC correlation (neuron) of SCKm and DistKm ROI type at each ROI count. (right vertical axis) The dashed line shows FC-SC detection (neuron) of SCKm and DistKm at each ROI count. **e,** Swarm plot of ROI volume of SCKm and DistKm at each ROI count.

**Fig. 3.**
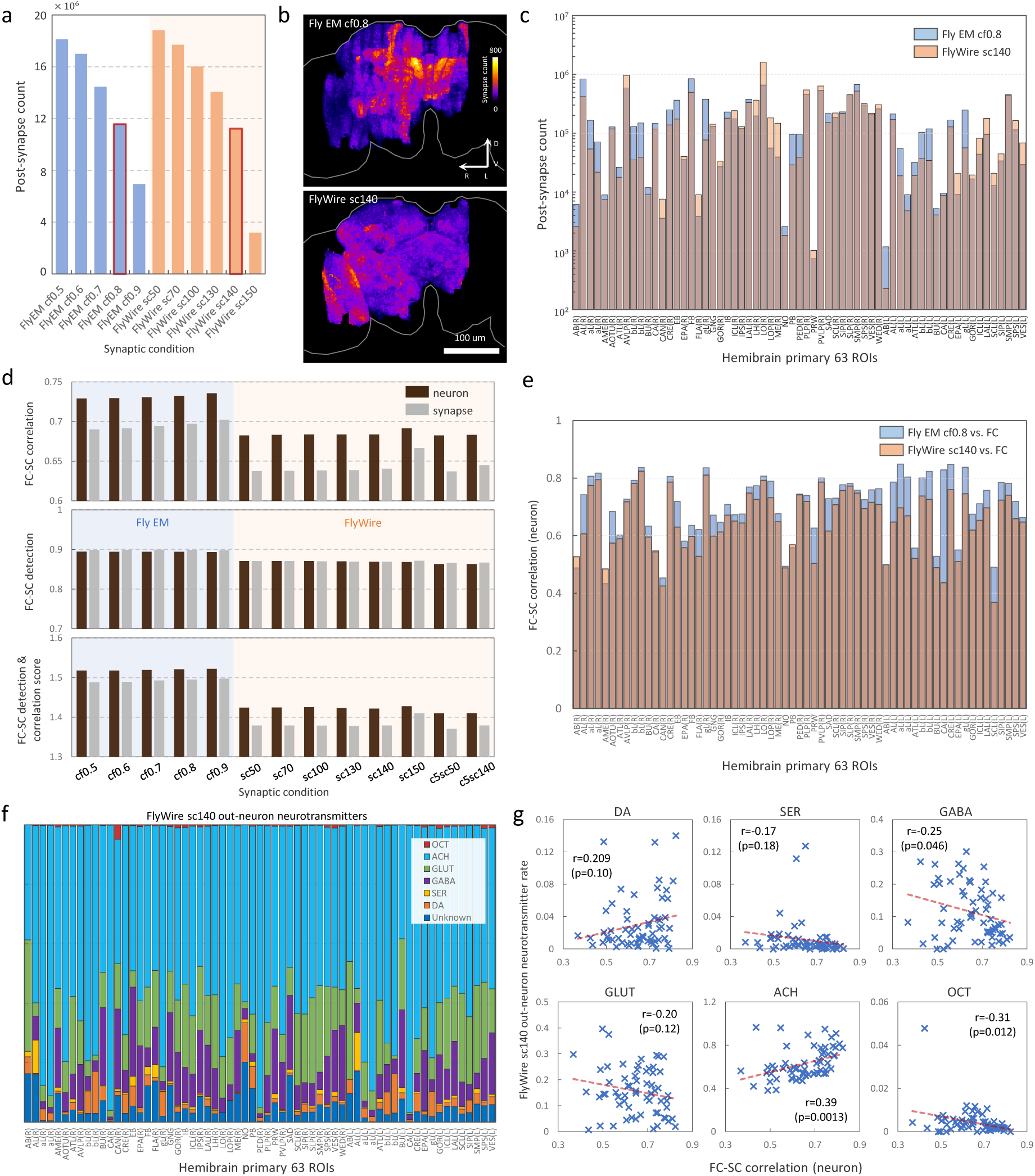
Comparison between FlyEM and FlyWire connectome data. a, Bar graph of post-synapse count in hemibrain region from FlyEM and FlyWire connectome data. The vertical axis shows post-synapse count, and the horizontal axis shows confidence thresholds for FlyEM and Cleft score thresholds for FlyWire. **b,** (top) Maximum Z-projection of synapse point cloud of FlyEM (confidence thresholds=0.8 [hereafter, cf0.8]) (bottom) Maximum Z-projection of synapse point cloud of FlyWire (Cleft score threshold=140 [hereafter, sc140]) **c,** Bar graph of post-synapse count in hemibrain primary ROIs. Blue bar shows FlyEM (cf0.8), red bar shows FlyWire (sc140). The vertical axis shows post-synapse count, and the horizontal axis shows hemibrain primary ROIs. **d,** Bar graphs of FC-SC correlation, FC-SC detection, and FC-SC Detection & Correlation score, respectively. The horizontal axis shows confidence thresholds for FlyEM and Cleft score thresholds for FlyWire. **e,** Bar graph of FC-SC correlation in hemibrain primary ROIs. Blue bars show FlyEM (cf0.8) vs FC (neuron), red bars show FlyWire (sc140) vs FC (neuron). The vertical axis shows FC-SC correlation, and the horizontal axis shows hemibrain primary ROIs. **f,** Bar graph of neurotransmitter rate of output neurons in hemibrain primary ROIs based on FlyWire (sc140) connectome data **[46]**. The horizontal axis shows hemibrain primary ROIs. (DA: dopamine, SER: serotonin, GABA, GLUT: glutamine, ACH: acetylcholine, OCT: octopamine) **g,** Scatter plots of neurotransmitter rate of output neurons vs. FC-SC correlation (neuron) in hemibrain primary ROIs (FlyWire sc140). Each of the six neurotransmitters was compared.

**Fig. 4.**
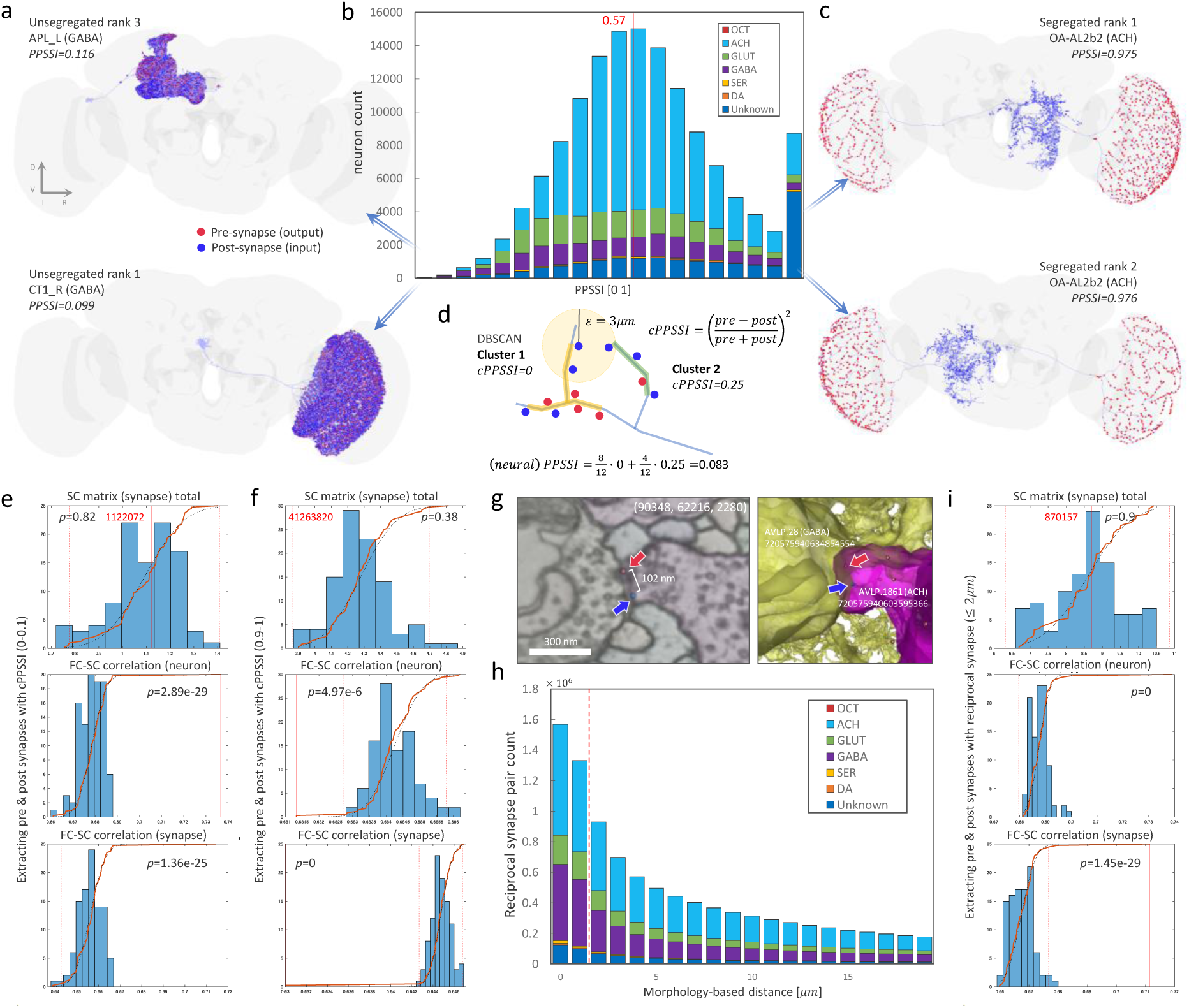
Quantification of pre-and post-synapse segregation, and the relationship between FC and synapses. a, Unsegregated first (CT1_R) and third (APL_L) ranked cells in the FlyWire (sc140) connectome data. A red dot shows pre-synapse, a blue dot shows post-synapse. **b,** The PPSSI histogram of all neurons (139,255) in FlyWire (sc140) connectome data. The vertical axis shows neuron count, and the horizontal axis shows PPSSI [0 1]. **c,** Segregated first (OA-AL2b2 cell) and second (OA-AL2b2 cell) ranking in FlyWire (sc140) connectome data. **d,** Schematic image of pre-and post-synapse clustering based on DBSCAN with a morphology-based distance threshold of 3 μm, and calculation of cPPSSI and PPSSI. **e,** Histogram of null SC matrices (blue bar) and extracted SC matrix with cPPSSI (0-0.1) (red solid line). Black dotted line shows cumulative distribution function of the normal distribution, brown solid line shows cumulative distribution function of null & extracted SC matrices, and red dotted line shows Bonferroni-corrected p<0.05 threshold. Top shows SC matrix (synapse) total, middle shows FC-SC correlation (neuron), and bottom shows FC-SC correlation (synapse). **f,** Histogram of null SC matrices and extracted SC matrix with cPPSSI (0.9-1). **g,** Example of a reciprocal-synapse in the EM image (left) and in 3D (right). **h,** Histogram of reciprocal-synaptic minimum distances in FlyWire (sc140) connectome data. The vertical axis shows reciprocal-synapse pair count, and the horizontal axis shows distance [*μm*]. **i,** Histogram of null SC matrices and extracted SC matrix for reciprocal-synapses (≤ 2*μm*).

## Results

### Investigation of the optimal nuisance factor removal method for calcium imaging

First, the optimal pre-processing of the calcium imaging data of *D*. *melanogaster* was investigated. Whole-brain calcium imaging data of *D*. *melanogaster* were acquired by Brezovec et al. [14] and are publicly available as an open dataset. We downloaded and used it for our study. The registration method used in fMRI was applied to the calcium imaging data (details in Methods, Ext.Data.Fig.1-1) to register to the FDA template [14] (Fig. 1b). Then, various nuisance factor removal methods (details in Methods) were applied to the ROI time-series to generate the FC matrices of eight flies (Fig. 1a). Three types of ROIs were used in the analysis: ROIs based on the anatomical neuropil, commonly used in *Drosophila* (hemibrain ROI) [42], ROIs based on k-means clustering of the SC matrix of 146,604 voxels in the hemibrain (SCKm), and ROIs based on distance-based clustering of voxels in the hemibrain (DistKm). SCKm can provide ROIs close to the shape of the anatomical neuropil across various ROI counts (Ext.Data.Fig.2-1a), while DistKm can provide ROIs that are completely unrelated to the anatomical neuropil. We investigated whether nuisance factor removal methods remained robust across these various ROI types. Next, a correlation analysis and ROC curve analysis were performed using the SC matrices based on FlyEM and the FC matrices with these ROI types (Fig. 1a). Fig. 1d-f shows the FC-SC correlation results, FC-SC detection results, and the normalized total score of FC-SC detection and correlation (details in Methods), respectively. Since signals are assumed to be transmitted between regions based on SC, when SC is treated as a biologically motivated validation metric, we considered a pipeline with higher similarity of FC-SC and higher detection to be better. After the comparison of the raw (leftmost) and removal methods for various ROI types, the combination of polynomial detrending [43] & tCompCor [44] was found to yield the optimal results. The optimal removal methods were found to differ between fMRI [10] and calcium imaging data, and these results are further described in the Discussion section.

### Comparison of the FC-SC correlation between humans and flies

Spatial smoothing is useful for absorbing inter-individual variability and conducting second-level group analysis [10, 16]. Here, we explored the effect of spatial smoothing (of the calcium imaging data) on various ROI counts in DistKm (Fig. 2a) and SCKm (Ext.Data.Fig.2-1a) ROIs. In DistKm, smoothing was highly effective, and the optimal kernel radius of the Gaussian filter was around 23 voxels (52.44 μm) for various ROI counts (Fig. 2b). In SCKm, smoothing was ineffective for ROI count below 100, while the kernel radius above 16 voxels (36.48 μm) was effective for ROI count above 100 (Ext.Data.Fig.2-1b). Increasing the smoothing size improved the correlation and AUC between group-averaged FC and SC, indicating the presence of inter-individual variability in FC. SCKm has a larger variation in volume for each ROI count than DistKm (Fig. 1c, Fig. 2e); as such, the effect of spatial smoothing is likely unstable. Next, we investigated whether the combination of spatial smoothing and Poly-tCompCor was effective. We found that this combination was significantly more effective than smoothing alone for SCKm and DistKm (Fig. 2c, Mann–Whitney U test p<1.67e-4 Bonferroni corrected). Conversely, the combination of a high-pass filter and nuisance factor removal overlapped for the purpose of noise removal, and the effect was degraded (Ext.Data.Fig.2-1d), as found in fMRI [10].

The optimal FC-SC correlation for various ROI counts was determined. Results showed that the correlation decreased proportionally to the logarithm of the ROI count (Fig. 2d). This was also compared between flies and mammals. At approximately 50 ROIs, the difference was nearly 1.08-fold to 1.22-fold (r=0.66 for humans and r=0.71-0.81 for flies [SCKm, DistKm]); and at approximately 1,000 ROIs, the difference was roughly 1.12-fold to 1.24-fold (r=0.54 for humans and r=0.6-0.67 for flies), which was not a considerable difference (not significant by Mann-Whitney U test, p=0.267). The difference was slightly larger (1.21 to 1.5-fold: r=0.378 for marmosets and r=0.46-0.57 for flies) than the marmoset results at around 10,000 ROIs (Ext.Data.Fig.2-1c). Although SCs for flies and humans were obtained from single individuals, the tracer data is a composite from 52 marmosets [45], which might contribute to the difference with the fly’s FC-SC correlation. In addition to log transformation, we calculated FC-SC correlation using Gaussian resampling [6, 7]. When the ROI count was small (200 or less), Gaussian resampling tended to produce a higher SC-FC correlation (Ext.Data.Fig.2-2a). As the ROI count increased so too did the Sparsity Rate (Ext.Data.Fig.2-2b) and log transformation tended to produce a higher SC-FC correlation (Ext.Data.Fig.2-2a). In both methods we still observed a tendency for the FC-SC correlation to decrease as the ROI count increased.

### Comparison between FlyEM and FlyWire connectome data

We used the FlyEM’s SC to compare the averaged FC of eight flies. In this section, we also used the FlyWire connectome data [46] to compare the FC and each SC, and possibly reveal the inter-individual variability of the two EM connectome data. As mentioned earlier, FlyEM has a synaptic “confidence” threshold and contains 21.4 million post-synapses in the hemibrain region at its initial value of 0. Conversely, FlyWire has a ‘CleftScore’ [19] threshold and 18.8 million post-synapses at its initial value of 50 in hemibrain primary ROIs. Since the number of post-synapses differs at each threshold, we set the confidence threshold of 0.8 as high as possible, thereby increasing the FC-SC correlation (Fig. 3d, ExtFig.3-1b). Then we adjusted the CleftScore to 140, which has a post-synapse number of approximately 11.4 million (Fig. 3a). These thresholds were used for all analyses in this study. When we displayed the maximum Z-projection of the synapse point cloud of post-synapse counts in each voxel, we found that the trends in synapse detection completely varied (Fig. 3b). In each region, FlyEM had more synapses in the mushroom bodies (aL, bL, etc.) and in the major neuropils of the central nervous system (CNS; EB, FB, NO, and PB). In comparison, FlyWire had more synapses in the visual system (LO and ME) and in the periesophageal neuropils (CAN and FLA; Fig. 3c). In the FC-SC correlation of each region, FlyEM had higher values in most regions at 58:5 (Fig. 3e, ExtFig.3-1a). As a result, FlyEM r=0.73 differed from FlyWire r=0.68 at the matrix level (Fig. 3d). Both datasets were from genotype Canton-S-G1×w1118 adult females; FlyEM was five-day-old [12], and FlyWire was seven-day-old [13]. Although brain development may differ depending on the rearing environment, the tendency to detect synapses substantially varied. This may be caused by differences in detection accuracy resulting from the resolution of EM scanning, but not to inter-individual variability.

We next used the FlyWire connectome data to investigate the ratios of neurotransmitter input (ExtFig.3-2a) and output (Fig. 3f) in each region. The highest percentage was acetylcholine (62.8%±14.2%), followed by glutamine (15.5%±9.2%), then GABA (11.4%±8.2%) in neurotransmitter output. The analysis of the relationship between this ratio and FC-SC correlation in each neurotransmitter revealed significant correlations for acetylcholine (r=0.39, p=0.0013) and GABA (r=-0.25, p=0.046) (Fig. 3g). That is, the higher the percentage of excitatory connections, the higher the FC-SC correlation; conversely, the higher the percentage of inhibitory connections, the lower the FC-SC correlation. In humans, FC-SC correlations are lower in the evolutionarily expanded Brodmann area 39,45 and the temporal lobe [8]; one possibility is that a high number of long-range inhibitory connections [47, 48, 49] may result in a low FC-SC correlation.

### Unsegregated synapses contribute considerably to functional connectivity

So far, we have compared FC and SC at the macroscopic region level. We next investigated the relationship between the synapses, the source of functional activity (pre-synaptic population dynamics), and FC. We examined the synaptic segregation in FlyWire neurons because many neurons showed pre-and post-synaptic intermingling. Then, we determined how unsegregated and segregated synapses affected the FC. To quantify synaptic segregation, we developed the PPSSI (Fig. 4d). Using DBSCAN [50] with a morphology-based distance threshold of 3 μm (details on Methods), we clustered synapses in each neuron and calculated the degree of segregation for each cluster (cPPSSI). This cPPSSI is also assigned to synapses in each cluster. PPSSI is then the average of the cPPSSI values of all synapses in a neuron. A similar index to our PPSSI, namely, the segregation index, is defined in the literature [22], but we have simplified from this. Since the cPPSSI of each synaptic cluster and the PPSSI of a cell are linearly related, they can be displayed as a common metric. Fig. 5h illustrates this, where the cPPSSI and PPSSI values within each synaptic cluster are uniformly displayed within the [0, 1] range. For clustering, we simply applied DBSCAN by using a morphology-based distance between synaptic nearest arbors. The segregation index is impossible to calculate for neurons such as AN and DN, which have only input or output synapses, because it results in calculations of 0 × ∞ or division by 0. Our coefficient simplifies the calculation and addresses these issues. Fig. 4b shows the result of applying PPSSI to 139,255 whole-brain neurons in the FlyWire dataset. This showed a histogram with the most frequent values near the center. When we created an unsegregated ranking (details in Methods) by considering the number of synapses, APL and CT1 cells occupied the top 1–4 positions, indicating that we efficiently extracted unsegregated neurons (Fig. 4a, Ext.Data.Fig.4-3a). APL was also at the top for FlyEM (Ext.Data.Fig.4-1a). Similarly, we established a segregated ranking and successfully extracted neurons with a high degree of segregation (Fig. 4c, Ext.Data.Fig.4-3; Ext.Data.Fig.4-1c and Ext.Data.Fig.4-4 for FlyEM).

**Fig. 5.**
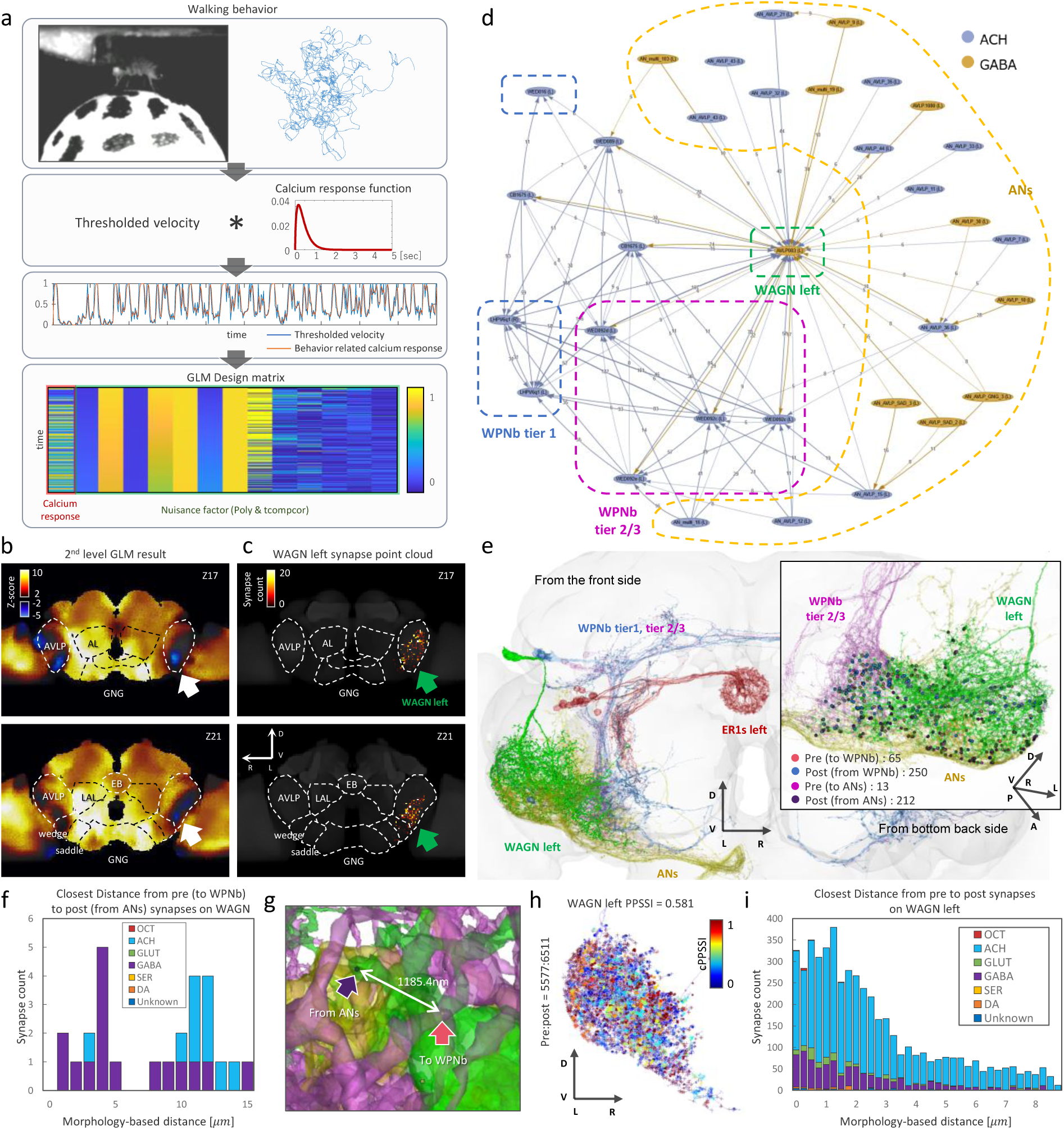
General Linear Model analysis result and Wedge-AVLP GABA neuron. a, Schematic image of GLM analysis. From the top, experimental image and walking trajectory, calcium response function, thresholded velocity, and GLM design matrix, respectively. **b,** 2^nd^ level (group) GLM analysis result from 9 flies. Top shows Z=17 slice of the resized FDA template, and the bottom shows the Z=21 slice. The white arrow indicates the suppressed region during the fly walking task. **c,** Synapse point cloud of the left WAGN in FlyWire (sc140) connectome data. Top shows Z=17 slice of the resized FDA template, and bottom shows the Z=21 slice. **d,** Network graph of the left WAGN, ANs, WPNb tier 1 and tier 2/3 neurons from FlyWire codex (sc50, connection threshold>5). Blue is acetylcholine, ocher is GABA neuron. The small number on the edges shows synapse count. **e,** 3D image of the left WAGN, ANs, WPNb tier 1, tier 2/3 and left ER1 neurons, frontal view. Inside the black square shows the left WAGN, ANs and WPNb tier 2/3 from the back side. **f,** Histogram of closest distance from pre-synapse (to WPNb) to post-synapse (from ANs) in the left WAGN. Horizontal axis shows morphology-based distance [*μm*]. Neurotransmitter indicates input ANs to post-synapses. **g,** Example 3D image of pre-synapse (to WPNb) to post-synapse (from ANs). **h,** All pre-and post-synapses of the left WAGN from the front. Color indicates cPPSSI in each synapse. **i,** Histogram of closest distance from pre-synapse to post-synapse in the left WAGN. The horizontal axis shows morphology-based distance [*μm*]. Neurotransmitter indicates input neurons to post-synapses.

Next, synapses with cPPSSI values between 0 and 0.1 were extracted, and the SC matrices (unsegregated synapses) of neurons and synapses were created. Synapses were then randomly sampled to generate 99 null SC matrices (random synapses), centered around the total number of connections (1,122,072) in the SC matrix (unsegregated synapses), forming a normal distribution (Fig. 4e, p=0.82, details in Methods). The FC-SC correlations of the SC matrices (unsegregated synapses) of neurons and synapses were significantly higher than those of the randomly extracted samples (Fig. 4e, p=2.89e-29, 1.36e-25; Ext.Data.Fig.4-1d for FlyEM). Therefore, unsegregated synapses might strongly contribute to FC. A similar analysis was performed on synapses with a cPPSSI of 0.9–1, which surprisingly showed a significantly lower FC-SC correlation (Fig. 4f, p=4.97e-6, 0; Ext.Data.Fig.4-1e for FlyEM). Therefore, segregated synapses may not be able to contribute to strongly to the FC, and these findings are further explored in the Discussion section.

Among unsegregated synapses, reciprocal synapses (Fig. 4g, Ext.Data.Fig.4-1f) have been well studied [33, 34, 39, 40, 41] and their function is considered to be clearer than unsegregated synapses. The shortest distance of such reciprocal synapses was extracted from whole-brain neurons, shown in Fig. 4h (Ext.Data.Fig.4-1g for FlyEM). They are particularly abundant within 2 μm (Ext.Data.Fig.4-2a,c for CT1 and APL cells), and acetylcholine and GABA constitute the majority (Fig. 4h). The comparison of the SC matrix based on reciprocal synapses (<2 μm) with the null SC matrices showed that the FC-SC correlation was significantly higher (Fig. 4I, p=0, 2.7e-32; Ext.Data.Fig.4-1h for FlyEM). Therefore, unsegregated synapses and reciprocal synapses (<2 μm) strongly contributed to FC. Furthermore, since the FC was the average of eight flies, the contribution of unsegregated synapses to FC may be a common phenomenon across individuals.

### Region suppression analysis during fly walking

The whole-brain calcium imaging dataset from Brezovec et al. [14] provides fly walking data. Since fMRI pre-processing methods can be applied to calcium imaging data, we investigated whether voxel-level analysis using a general linear model (GLM) [51] could be applied to the fly walking task (Fig. 5a, details in Methods). Fig. 5b shows the GLM analysis result. Similar to the results of Brezovec et al., our findings revealed that LAL, FB, and IPS neuropils showed strong activity; the GNG neuropil, which is highly connected to leg neuropils in the ventral nerve cord [52], also exhibited strong activity. Although Brezovec et al. detected voxels that are strongly correlated with velocity, the GLM extracted voxels associated with walking behavior. Moreover, we found inactivity in the Wedge and AVLP neuropils. We searched for a GABA neuron that would fit with this phenomenon and managed to find one that extended its dendrites widely from the Wedge to AVLP neuropils (Fig. 5e, h, Root ID: 720575940644632087). The synapse point cloud could also fit in this region (Fig. 5c). We called this neuron the Wedge-AVLP GABA neuron (WAGN). The Wedge neuropil receives mechanosensory (wind, sound) input and connects to WPNb neurons (Ext.Data.Fig.5-1a,b), which then connect to ER1 ring neurons that act as the fly’s compass [53, 54]. Interestingly, the left WAGN was strongly interconnected with WPNb tier 2/3 (left) [55] and received input from 22 ascending neurons (ANs; Fig. 5d). Therefore, it may be a pathway that provides walking behavior feedback to the compass; however, wind caused by walking was absent in the experimental setup (the fly was fixed and walked on a treadmill), and may relate to the strong regional suppression that was observed. As such, further experimentation is needed to pursue this topic.

The left WAGN resembles a hub neuron; however, it contains many unsegregated synapses (PPSSI=0.581). Segregated and unsegregated synapses are mixed throughout the cell, which is reflected in the intermediate PPSSI values (Fig. 5h). It received input from ANs (212 synapses) mainly on the ventral side, and outputs to WPNb (65 synapses), widely from the medial to the lateral side (Fig. 5e). A histogram of the shortest morphology-based distances between pre-synapses to WPNb and post-synapses from ANs is shown in Fig. 5f, and the closest distance was 1185.4 nm (Fig. 5g). Many acetylcholine inputs (disynaptic inhibition [28]) were closer than this value (Fig. 5i), and many GABA inputs from ANs (Fig. 5f) further suppress such disynaptic inhibition. AVLP ventral suppression (Fig. 5b white arrows) may be the suppression from these ANs. Thus, a highly delicate and localized control is possibly exercised from WAGN to WPNb tier 2/3 (left).

## Discussion

Although nuisance factor removal methods such as global mean (GM) and aCompCor [44] are optimal for fMRI [10], calcium imaging showed different results. GM is the average of all voxels, and the regression of this signal is useful in fMRI because noise from the equipment and coils affects the entire voxel image. Conversely, in two-photon microscopy, noise does not affect all voxels because of the excitation of specific molecules by laser absorption, and GM does not show a positive effect. aCompCor extracts the component time-series from the voxel time-series of white matter and cerebrospinal fluid (CSF) via PCA and then performs a regression. In flies, the CSF had no ROI mask, but fibers had ROIs; therefore, fibers were used instead of the white matter. However, many synapses may exist around these fibers, and aCompCor was not as effective as in humans. Conversely, polynomial detrending [43] was highly effective (Fig. 1d-f), likely because of linear thermal noise and other factors caused by the continuous excitation of molecules (for 30 min). tCompCor [44] is the component time-series extracted by PCA from voxels with the top 1000 standard deviations among all voxels. It potentially captures minute fluctuations caused by fly movements and effectively removes them.

The SC matrix from a cPPSSI of 0.9–1 synapses resulted in a significantly lower FC-SC correlation than the SC matrix from randomly selected synapses (Fig. 4f for FlyWire, Ext.Data.Fig.4-1e for FlyEM). This result was surprising, but we propose several possible reasons. Neurons with highly segregated synapses include ascending neurons (ANs) and descending neurons (DNs) (Ext.Data.Fig.4-3b). They also have a small number of unsegregated synapses (i.e. Sep rank11: PPSSI=0.963). They are thought to be specialized for input or output in the CNS regions and may show large differences between SC and FC. Another reason is that these neurons may have widespread logical OR inputs and outputs. In Fig. 4c, an action potential at one point in the input synapse (blue) is likely transmitted to all red synapses. Such an OR circuit-like disproportionate input–output possibly causes a mismatch with the FC, because the functional activity in the output region originates from activity at a single location. These neurons are likely those whose FC are difficult to capture. Conversely, neurons with highly unsegregated synapses may capture the inputs of neighboring synapses, such as forming a logical AND circuit, correlating well with the output regions.

The FC-SC correlation was highly dependent on ROI count in both flies and humans (Fig. 2d). Through optimal nuisance factor removal and smoothing, we achieved r=0.87 at 20 ROIs. Furthermore, by applying Spearman correlation instead of Pearson, we obtained r=0.928. Thus, for a reasonably small ROI count the correlation was very high; however, for 10,000 ROIs, the correlation decreased to r=0.57. This phenomenon could be attributed to the presence of highly segregated synapses. Such neurons are widespread in the CNS (Fig. 4b, Ext.Data.Fig.4-2f) and are thought to provide logical OR circuits. When ROI volume was large (when the ROI count was small), a single ROI could cover the distributed input synapses. However, when a ROI was divided, input synapses were also divided across many ROIs, and the mismatch between SC and FC would increase. Other possibilities include the presence of gap junctions and FC due to neuropeptides [56], but they are outside of the scope of the present study and their relationship with ROI count is unclear.

In this study, we applied fMRI techniques directly to calcium imaging in flies and our results suggest many similarities between humans and fly neural computation (Fig. 2d). The calcium signal in the fly brain represents pre-synaptic population dynamics and the BOLD (Blood Oxygenation Level Dependent) signal in the mammalian brain is also thought to contain the same type of population dynamics. It has been estimated that 43% of the gray matter’s energy consumption is due to synaptic activity [57]. Local oscillations of synaptic activity are detected in the BOLD signal in fMRI through neurovascular coupling [58]. However, the BOLD signal contains biological noise such as cardiac or respiratory noise, etc. We used nuisance factor removal methods to remove these noise sources to approximate pre-synaptic population dynamics and calculate inter-regional FC. Despite the differences in modalities, this commonality in FC is likely responsible for the observed trends in high FC-SC correlation. Studies on FC-SC correlation have also been conducted in C. elegans [59, 56], but results showed a low correlation. In C. elegans, the FC of calcium dynamics in the soma was compared with the SC of the whole cell. Neurons in C. elegans are non-spiking, and calcium dynamics have been shown to differ between compartments in a neuron [60]. While fly and human studies compared the SC and FC of pre-synaptic population dynamics in brain regions, C. elegans studies compared the SC of cells (not regions) with the FC of dynamics in the cell body (in terms of compartments). This discrepancy is thought to explain the differing results. For this study, we used FC-SC correspondence as a biologically motivated validation metric for investigating nuisance removal methods. Numerous studies have demonstrated statistically significant FC-SC relationships across species and modalities (as presented in our Introduction). These findings suggest that SC provides an important anatomical constraint on FC and we were able to obtain reasonable results based on our approach. We recognize that biologically meaningful functional relationships may also arise through indirect pathways, network effects, neuromodulation, and other mechanisms not captured by direct SC. Although SC cannot be equated with FC, we believe that the relationship between the two remains strong. To investigate this matter further, we consider it necessary to conduct additional research involving the simultaneous measurement of whole-brain FC (calcium) and ‘non-synaptic’ signals.

Our study revealed that unsegregated synapses [28, 33, 34, 39, 40] are widespread in neurons throughout the brain (Fig. 4b, Ext.Data.Fig.4-2f). They are evenly distributed across cells of various neurotransmitters, and some may be non-spiking [24, 25, 26, 27, 28, 29]. Retinal amacrine cells have extensive reciprocal synapses [33, 34, 39, 40, 41], which provide local feedback inhibition of burst inputs to improve signal dynamic range [34]. Grimes et al. [34] noted “The retina is a beautiful example of a neural network that optimizes signal processing capacity while minimizing cellular cost.” To maintain signal dynamic range, A17 amacrine cells must optimize processing units and wiring costs. If one unit equaled one cell, an enormous number of cell bodies would be required, reducing the number of processing units per volume and increasing the energy cost during development. To optimize this, Grimes et al. proposed arranging units capable of parallel processing within a single cell, thereby maximizing processing unit count and wiring cost per volume. Signal bursts might also occur in the CNS, in which case CNS neurons would also require dynamic range adjustment. The idea of optimizing processing unit count per volume is highly compelling and is thought to apply not only to the retina but throughout the entire brain. This design concept may therefore be useful for studies across multiple species.

## Materials and Methods

### Preprocessing of *D. melanogaster* calcium imaging data

Whole-brain calcium imaging data of *D*. *melanogaster* were acquired by Brezovec et al. [14] and downloaded from DANDI (https://dandiarchive.org/dandiset/000727/0.240106.0043). 4D NIfTI image data (256×128×49 voxels, 3384 frames) were extracted from a NWB file by an in-house MATLAB script. The FDA template, made from in-vivo calcium imaging, was resized (256×128×49 voxels, 2.45×2.28×3.72*μ*m for a voxel) and used as a template because the original thresholded FDA template [61] was extremely large (1652×768×479 voxels), and the calcium imaging file was large (more than 40 GB); therefore, a smaller template with the imaging data was preferably used. Preprocessing and image registration were performed using Statistical Parametric Mapping (SPM12) [15] and ANTs [62] (Ext.Data.Fig.1-1a). Motion and slice timing correction were performed using SPM12, and the averaged NIfTI image was registered by ANTs to the resized FDA template. Lastly, 4D NIfTI images were transformed using the transform information. The data contained n=9 flies, but 8 flies were used for the ROI-based analysis because of missing time-series in some ROIs in one fly. Nine flies were used in the GLM analysis.

Combinations of the following nuisance factor removal methods were investigated: global mean (average signal across all voxels), global signal (average signal across all brain voxels), mean white matter (average signal across fiber voxels), 6HMP (head motion parameters), 24HMP, aCompCor, tCompCor [44], and polynomial detrending (Poly) [43]. These methods were implemented by an in-house MATLAB script [63]. A full width at half maximum (FWHM) from 1 voxel (2.28 *μ*m) to 30 voxels (68.4 *μ*m), at a 1 voxel step, was investigated to determine the optimal spatial smoothing size for calcium imaging data. Temporal high-pass filters were also examined from 0.1 Hz to 0.001 Hz.

### Transformation of EM connectome data into the resized FDA template

In the FlyWire case, connectome data (v783), such as neurons and synapses, were downloaded from FlyWire Codex (https://codex.flywire.ai/) and Zenodo (https://zenodo.org/records/10676866) [13, 64, 46]. Detailed synapse data were saved in an Apache feather format, which was then converted to a CSV file. This dataset contains 139,255 neurons and 54.5M (pair of pre-and post-) synapses as described in Dorkenwald et al. [13], with 18.8M post-synapses in the regions corresponding to the hemibrain primary ROIs. Synapse points were transformed from the FlyWire space to a JRC2018 female template [65] by the Python navis-flybrains package [65]. Since FlyWire coordinates are left and right reversed [13], they were flipped on the JRC2018 female template, and these points were transformed to the resized FDA template by ANTs. The use of these two templates facilitated precise data alignment across different modalities.

In the FlyEM case, the latest connectome data (v1.2) were downloaded from Janelia’s DVID website (https://dvid.io/blog/release-v1.2/). This data is provided in the format defined in https://neuprint.janelia.org/public/neuprintuserguide.pdf, and we extracted the neurons and synapses from it. The entire segmentation body is 28M segmentations, containing 99,644 Traced neurons. In addition, there were 73M (pre-or post-alone) synapses, 87M *synapseSets* records and 128M *synapseSet-to-Synapse* records. When we extracted post-synapses between Traced neurons, the total number was 21.4M (i.e., connections from Traced neurons to other body fragments, like Orphans, were excluded). Like those in the FlyWire case, synapse points were transformed from the FlyEM space to a JRC2018 female template by the Python navis-flybrains package. These points were transformed to a resized FDA template by ANTs.

### Generation of regions of interest for the hemibrain

The hemibrain neuropil atlas (114 ROIs, including 63 primary and 51 sub-regions, aligned with the FlyEM space) was downloaded from Virtual Fly Brain (https://www.virtualflybrain.org/). ROI files were transformed from the FlyEM space to a JRC2018 female template by Java template-building scripts (https://github.com/saalfeldlab/template-building) [65]. They were transformed to a resized FDA template by ANTs. In Fig. 3, 63 primary ROIs mentioned above were used. The 52 ROIs in Fig. 1 were selected from the primary ROIs, and approximately 50 ROIs (those used by Turner et al. [9]) were selected. ROIs were segmented by SCKm (Fig. 1 and Fig. 2) via the k-means clustering of the SC matrix of 146,604 voxels in the hemibrain. They were also segmented by DistKm (Fig. 1 and Fig. 2) via the k-means clustering of the distance of 146,604 voxels in the hemibrain.

The SC matrix was calculated according to synapse locations and ROIs. Since synapse data formats differ between FlyEM and FlyWire, they were standardized by using post-synapses within ROIs. The input and output neurons at each ROI were extracted from the paired pre-and post-synapse data. Then the input and output neurons and their synapses that were common between ROI pairs were counted to create SC matrices for neurons and synapses. The FC matrix was calculated using the pre-processed voxel time-series and ROIs. The intensity of the voxels in the ROI was averaged to obtain the average intensity time-series of the ROI. Then, Pearson correlation was calculated between ROI pairs to generate the FC matrix. The correlation coefficients were z-transformed, calculated for eight flies, and then averaged to obtain the final FC matrix.

### FC-SC detection and correlation

FC-SC correlation was calculated by taking the Pearson correlation between the logarithm of the SC matrix and the FC matrix, as done in other studies [6, 7, 10]. In the SC matrix, elements with zero connections become *-Inf* when logarithmically converted, so we substituted zero for them. Also, the diagonal elements of the FC matrix become *Inf* when z-transformed, so we replaced them with the maximum value of the matrix. For one FC matrix, we calculated its correlation with both the neuron SC matrix and the synapse SC matrix. If there is a neuron from post-synapse to pre-synapse between two regions, that neuron is counted as one, and the number of synapses in the connected region is used as the synapse SC matrix. FC-SC detection was calculated by ROC (Receiver Operating Characteristic) curve analysis. This gives the detection accuracy of the FC matrix when the SC matrix is used as ground truth. Thresholds were taken from 0% to 99% by 1% steps of all elements of the SC matrix, and 100 AUCs (Area Under the Curve) were calculated. The average of these AUCs is called the FC-SC detection. FC-SC correlation indicates the similarity of the two matrices, while FC-SC detection indicates the detection accuracy. Although the digit number of connections differs between neurons and synapses, FC-SC detection can absorb such differences. The FC-SC Detection & Correlation score was calculated as follows:

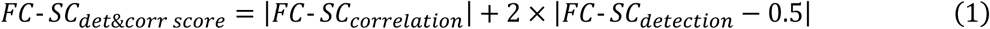

The FC-SC Detection & Correlation score is in the range of [0 2], where the larger the score, the higher the correlation and detection accuracy.

### Pre-and Post-synapse Segregation Index (PPSSI)

The PPSSI (Fig. 4d) was used in DBSCAN [50] with a distance threshold of 3 μm to generate pre-and post-synaptic clusters. This distance is not a straight-line distance, but rather a morphology-based distance between synapses along the nearest arbors (Ext.Data.Fig.4-1i). The threshold of 3 μm was empirically defined by several criteria. Firstly, CT1 and APL cells have the shortest distance of reciprocal synapses, within 2 μm (Ext.Data.Fig.4-2a,c). In zebrafish, reciprocal synapses were defined within 2.5 μm [41]. Schneider-Mizell et al. [22] defined a twig as a branch within 3 μm from the backbone of the cytoskeleton, and 80% of synaptic inputs in motor neurons are in the twig. In addition, 97% of pre-synaptic sites are located at less than 3 μm from the mitochondria. Therefore, 3 μm was deemed appropriate for the synapse cluster distance threshold. Once the synapse clusters were determined by DBSCAN, the degree of synaptic segregation for each cluster was calculated using the following formula:

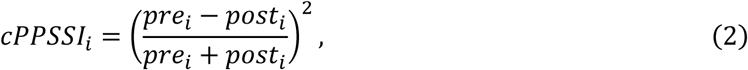

where *pre_i_* and *post_i_* are the pre-and post-synapse counts in cluster *i*, respectively. The cPPSSI of each cluster (each synapse) is in the range of [0 1]. The PPSSI of each neuron was calculated as follows:

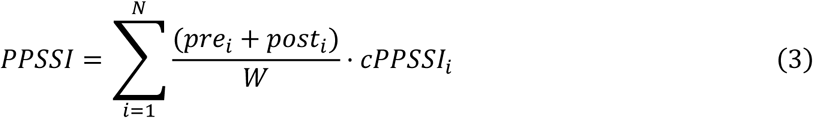

where *N* is the cluster number, and *W* is the total synapse count for a neuron. This cPPSSI is also assigned to synapses in each cluster. The PPSSI is then the average of the cPPSSI values of all synapses in a neuron.

The unsegregated ranking was calculated based on the PPSSI and total synapse count of neuron k:

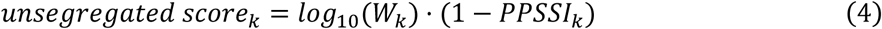

where *PPSSI_k_* is the PPSSI of neuron k and *W_k_* is the total synapse count of neuron k. This allows us to extract neurons with many inputs and outputs and highly unsegregated synapses. The segregated ranking was also calculated in the same way, using the following score:

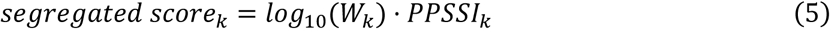

### GLM Analysis for the fly walking task

GLM analysis (Fig. 5a) was performed to investigate the whole-brain functional activity during the fly walking task. Movement velocity data were extracted from an NWB file by an in-house MATLAB script. Walking was defined as 1σ (34%) or more of the movement velocity and divided into walking and other. Less than 10% of movement velocity was zero cut. The canonical calcium response function (CRF) was created by the gamma probability density function (a=1.7, b=0.25) in MATLAB based on the GCaMP6f response [66]. The walking-task time-series was transformed into a calcium response time-series by convolution with the CRF. GLM design matrices were created using the calcium response time-series and nuisance factors (polynomial detrending and tCompCor). In whole-brain voxels, smoothing was applied with a kernel radius of 8 voxels (18.24 μm). Then, a Tukey taper (taper size = 8) was used for GLM pre-whitening [67]. A mixed-effects model [16] was used for group analysis, and the t-value of each voxel was calculated via the second-level analysis of OLS regression with a Tukey taper. These analyses were implemented using in-house MATLAB scripts [68].

## Statistical Analysis

A Mann–Whitney U test was used to examine whether the combination of spatial smoothing and nuisance factor removal was effective. Statistical significance was set at *p*<0.05, and Bonferroni correction was applied to correct the familywise error (FWE) rate. A permutation test was performed to analyze whether specific synapses (segregated, unsegregated, or reciprocal synapses) affect SC-FC correlation. The SC matrix (containing specific synapses) in the 63 ROIs in the hemibrain was generated, and its SC-FC correlation was calculated. Then, 99 null SC matrices (containing random synapses) were generated so that they would be normally distributed around the total number of connections in the SC matrix. Because the sparsity rate of the matrix also affects SC-FC correlation, we generated null SC matrices to match the sparsity rate of the target SC matrix (Ext. Data Fig.4-5). Since the SC-FC correlations of the null SC matrices also roughly followed a normal distribution, the T-value of the SC-FC correlation of the SC matrix was calculated using the cumulative distribution function of the normal distribution. For GLM analysis, a mixed-effects model was used for group analysis.

## Acknowledgements

The authors wish to thank Dr. Clandinin for providing valuable calcium imaging data and answering our question.

This research was supported by the program for Brain Mapping by Integrated Neurotechnologies for Disease Studies (Brain/MINDS 2.0) from the Japan Agency for Medical Research and Development, AMED. Grant number: JP24wm0625408.

## Data Availability

Whole-brain calcium imaging data and walking behavior data in *Drosophila Melanogaster* can be downloaded from the DANDI web site (https://dandiarchive.org/dandiset/000727/0.240106.0043). FlyWire connectome data can be downloaded from flywire codex (https://codex.flywire.ai/) and the Zenodo web site (https://zenodo.org/records/10676866). FlyEM connectome data can be downloaded from Janelia’s DVID web site (https://dvid.io/blog/release-v1.2/).

## Code Availability

Software and code used in this study is open source and available from https://github.com/takuto-okuno-riken/flywalk

## Authors’ Contributions

T.O. conceived of the presented idea. T.O. developed the theory, performed the computations. T.O. downloaded and processed fly data. T.O., A.W., H.O. and J.H. discussed the results and contributed to the final manuscript.

## Extended Data Figures

**Extended Data Fig.1-1.**
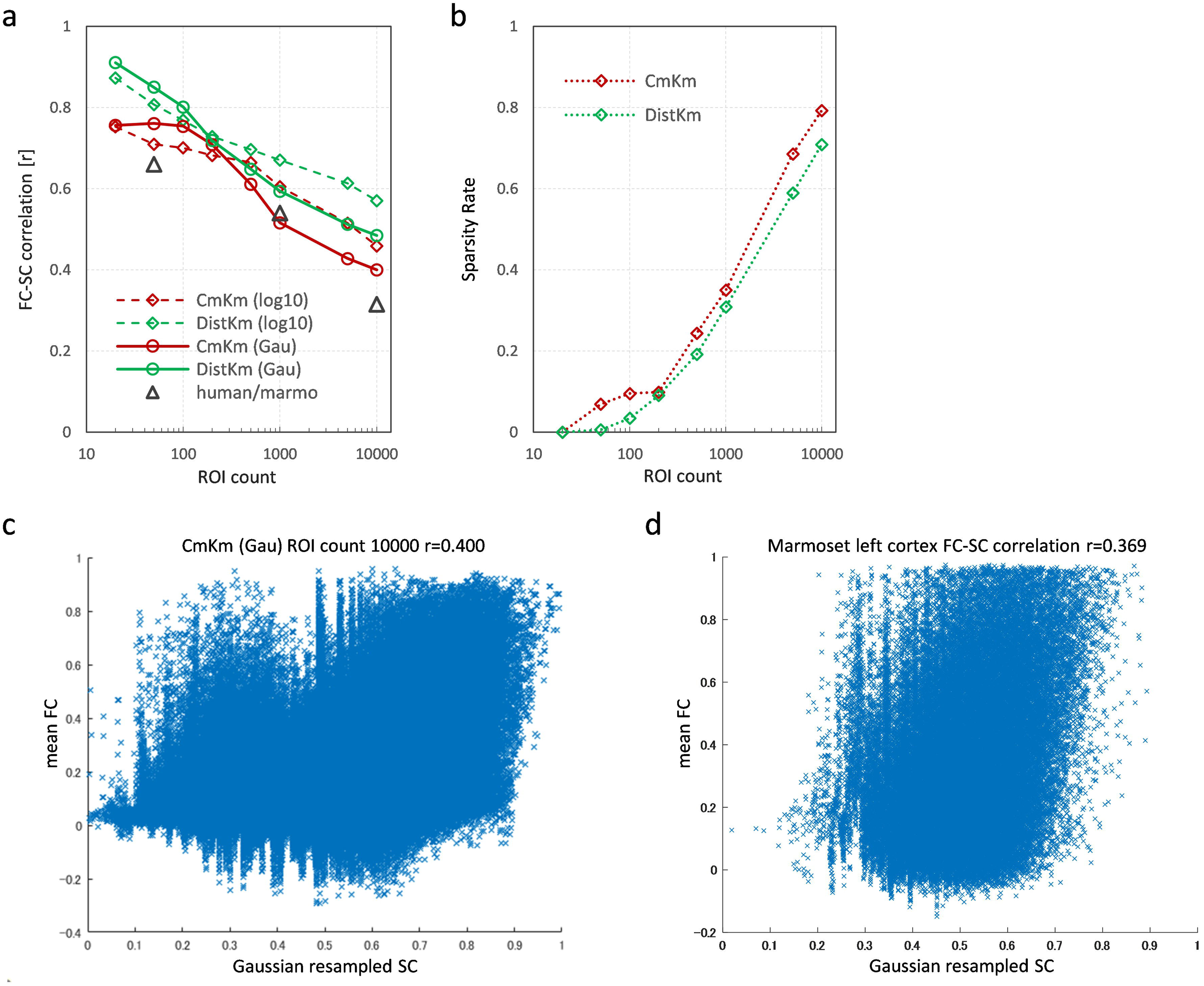
Schematic image of registration process and connectome data transformation in *Drosophila melanogaster.* a, Schematic image of calcium image registration to the resized FDA template. Motion and slice timing correction were conducted using SPM12 [15], and the averaged NIfTI image was registered by ANTs [62] to the resized FDA template. Lastly, 4D NIfTI images were transformed using the transformation information. **b,** Schematic image of synapse points transformation from FlyWire space to the resized FDA template. **c,** Schematic image of synapse points transformation from FlyEM space to resized FDA template.

**Extended Data Fig.1-2.**
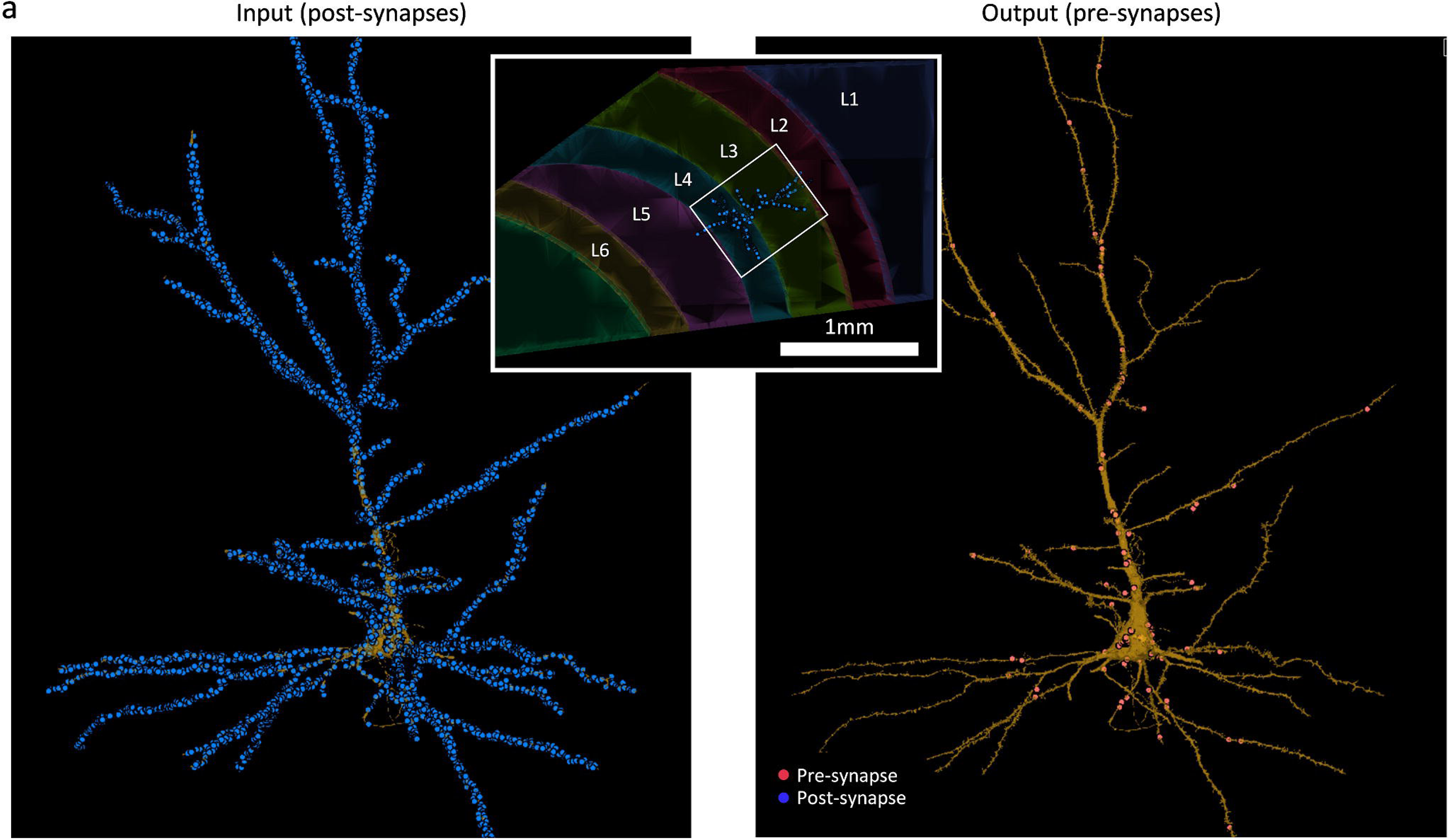
Pre-and Post-synapses of pyramidal cell in human temporal cortex layer 3-4. a, (left) Input to (post-synapses) a pyramidal cell in the human temporal cortex, from EM connectome data [38]. (center insert) A slice of human temporal cortex and target pyramidal cell. (right) Output from (pre-synapses) a pyramidal cell in the human temporal cortex, from EM connectome data. Blue dots show post-synaptic sites, Red dots show pre-synaptic sites.

**Extended Data Fig.2-1.**
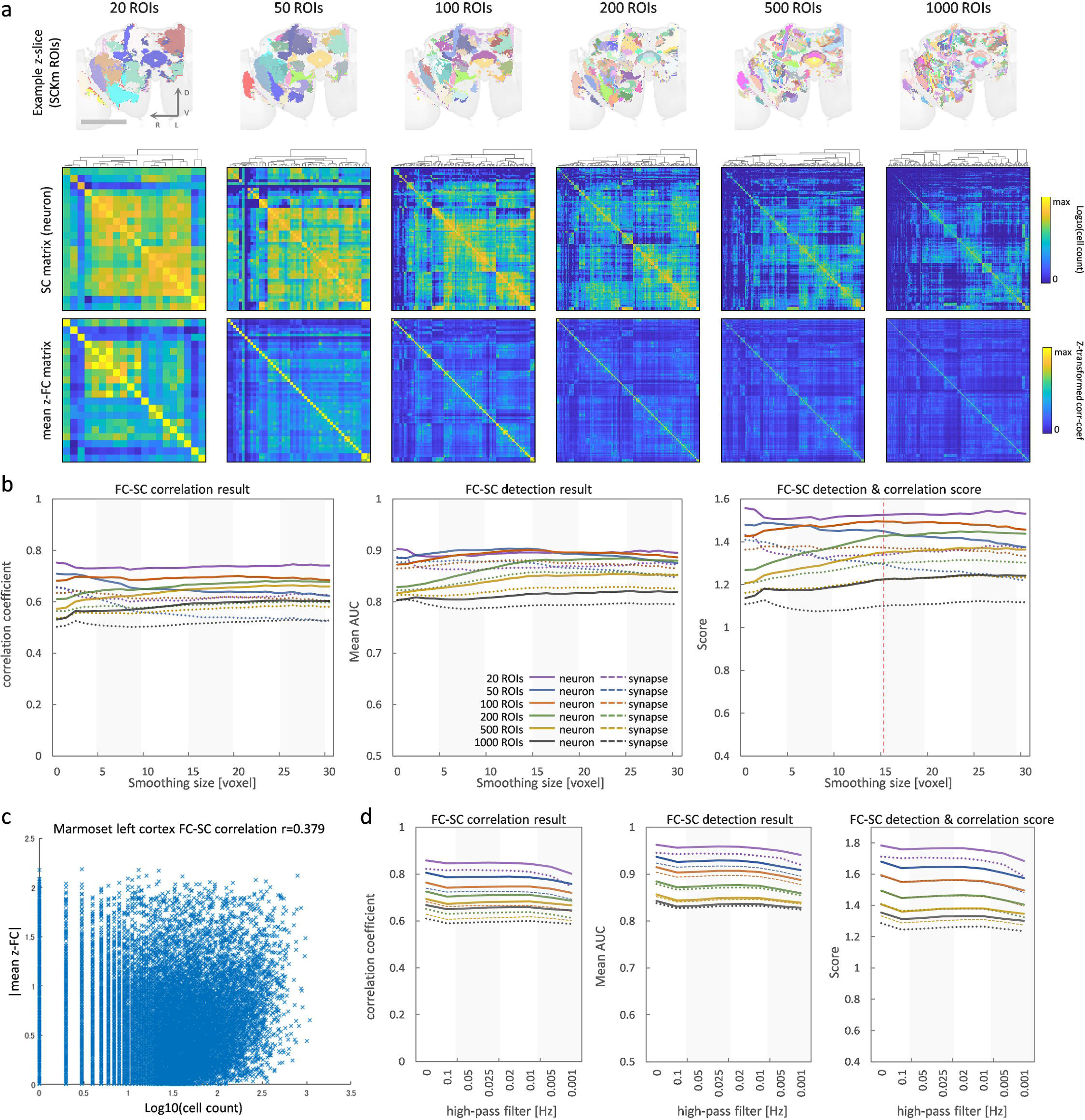
Investigating spatial smoothing size, and high-pass filtering. a, (top) Example z-slice of reduced FDA template with SCKm ROIs from 20 to 1000 ROIs. (middle) SC matrices (neuron) of SCKm ROIs from 20 to 1000 ROIs. (bottom) FC matrices of SCKm ROIs from 20 to 1000 ROIs. **b,** From left to right, FC-SC correlation, FC-SC detection, and FC-SC Detection & Correlation score, respectively. Solid lines shows FC vs. SC matrix (neuron), and dashed lines shows FC vs. SC matrix (synapse) for each ROI count. The vertical axis shows the correlation coefficient, averaged AUC, and FC-SC Detection & Correlation score, respectively. The horizontal axis shows spatial smoothing size (voxels). **c,** Scatter plot of FC-SC correlation in the marmoset left cortex [10] (145 injection×9862 voxels, r=0.379). **d,** Result of high-pass filtering and nuisance factor removal combination of DistKm ROIs. From left to right, FC-SC correlation, FC-SC detection, and FC-SC Detection & Correlation score, respectively. The high-pass filter degraded the FC-SC relationship.

**Extended Data Fig.2-2.**
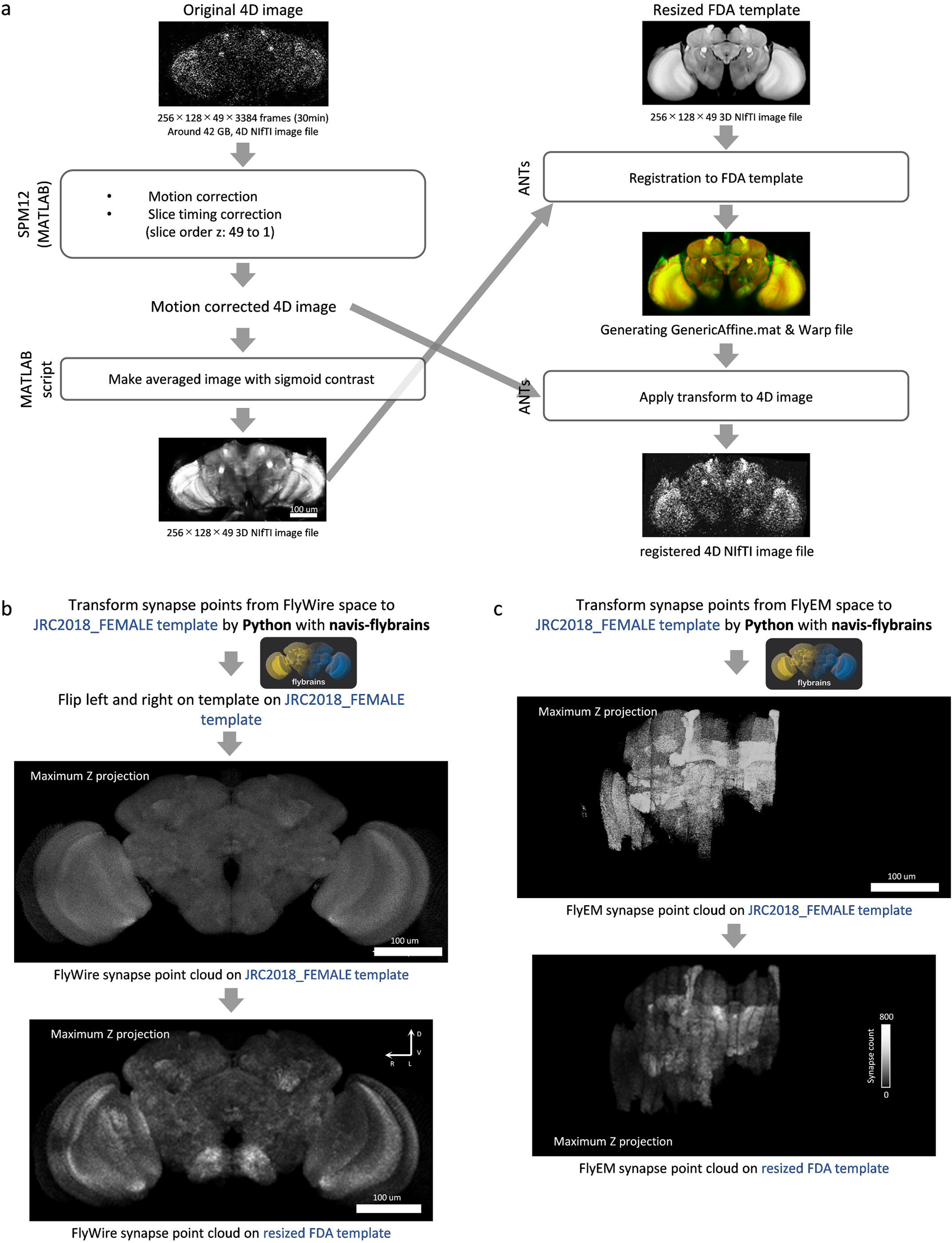
Investigating gaussian resampled SC vs. log-scaled SC. **a**, Relation between FC-SC correlation and ROI count. Solid line shows FC-SC correlation (neuron) of SCKm and DistKm ROI type with gaussian resampled SC. Dashed line shows FC-SC correlation with log-scaled SC. **b**, Sparsity rate by ROI count and ROI type. **c**, Example scatter plot of FC-SC correlation for a CmKm ROI count of 10000 with gaussian resampled SC. **d**, Scatter plot of FC-SC correlation in the marmoset left cortex with gaussian resampled SC.

**Extended Data Fig.3-1.**
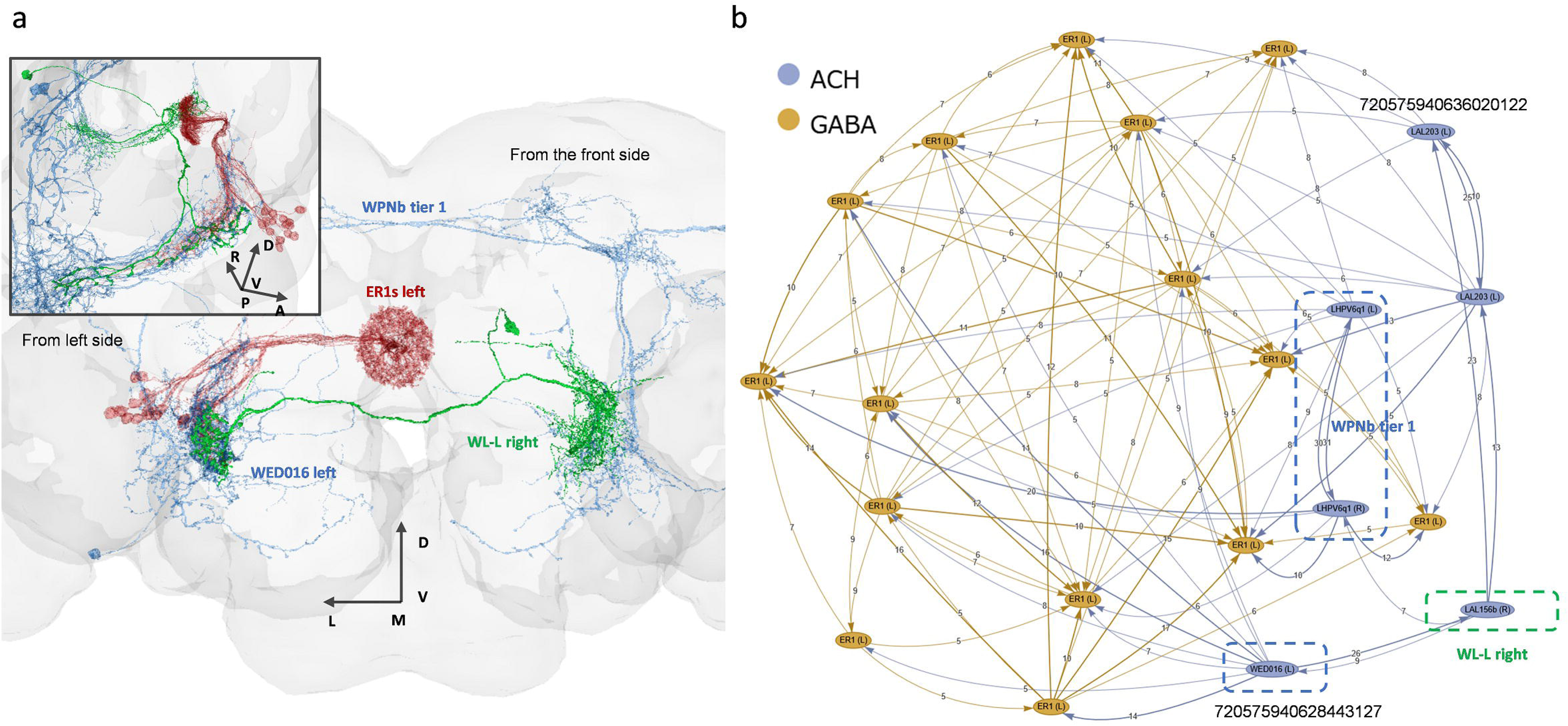
Comparison between FlyEM and FlyWire connectome data. a, (left) A scatter plot of regional post-synapse count between FlyEM (cf0.8) and FlyWire (sc140) in the hemibrain primary 63 ROIs. The Pearson correlation is not so similar, r=0.778, but the Spearman correlation is much more similar, r=0.9. (right) A scatter plot of log10 of the regional post-synapse count between FlyEM (cf0.8) and FlyWire (sc140) in the hemibrain primary ROIs. Pearson correlation shows r=0.919, a higher similarity. **b,** From left to right, FC-SC correlation, FC-SC detection, and FC-SC Detection & Correlation score, respectively. Solid lines shows FC vs. SC matrix (neuron), and dashed lines shows FC vs. SC matrix (synapse) of FlyEM (cf0.5-0.9) and FlyWire (sc50-140) corresponding Fig.3d. The vertical axis shows the correlation coefficient, averaged AUC, and FC-SC Detection & Correlation score, respectively. The horizontal axis shows spatial smoothing & nuisance factor removal combinations. **c,** From left to right, SC matrix (neuron) (FlyEM cf0.8), SC matrix (neuron) (FlyWire sc140) and FC matrix (Poly-tCompCor, no smoothing) in the hemibrain primary 63 ROIs. **d,** (right) SC matrix (neuron) result of FlyEM (cf0.8) minus FlyWire (sc140) in hemibrain primary ROIs. (left) SC matrix (synapse) result of FlyEM (cf0.8) minus FlyWire (sc140). FlyEM shows a higher number of synaptic connections.

**Extended Data Fig.3-2.**
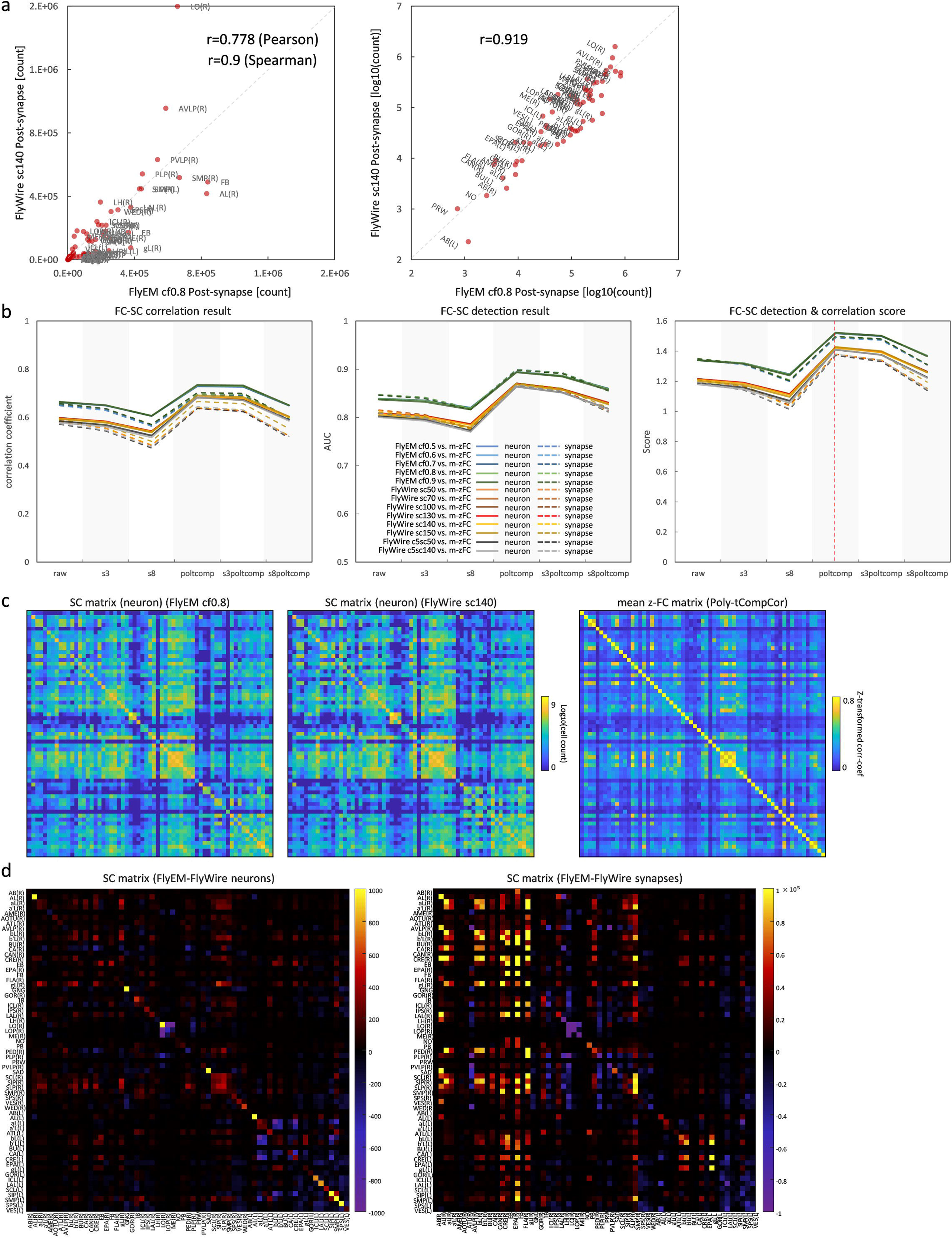
In-neuron neurotransmitters and comparison between FlyEM and FlyWire connectome data. a, Bar graph of neurotransmitter rate of input neurons in hemibrain primary 63 ROIs based on FlyWire (sc140) connectome data [46]. The horizontal axis shows hemibrain primary ROIs. (DA: dopamine, SER: serotonin, GABA, GLUT: glutamine, ACH: acetylcholine, OCT: octopamine) **b,** Scatter plots of neurotransmitter rate of input neurons vs. FC-SC correlation (neuron) in hemibrain primary ROIs (FlyWire sc140). Each of the six neurotransmitters was compared. **c,** Comparison between FlyEM and FlyWire APL right neuron (synapse point cloud mask). Red voxels are FlyEM, green voxels are FlyWire, and yellow voxels are both. Sørensen-Dice index (0.716) shows similarity, so neuropil shape is similar between two. **d,** (left) A scatter plot of post-synapse count between FlyEM and FlyWire APL right neuron. Pearson correlation shows r=0.104, so not similar at all. (center) Same as left, but synapse point cloud was spatially smoothed by gaussian kernel (2 voxels). Similarity was increased (r=0.379). (right) Same as center, but smoothed by gaussian kernel (8 voxels). Similarity was increased (r=0.66). These results support that spatial smoothing can absorb inter-individual variability.

**Extended Data Fig.4-1.**
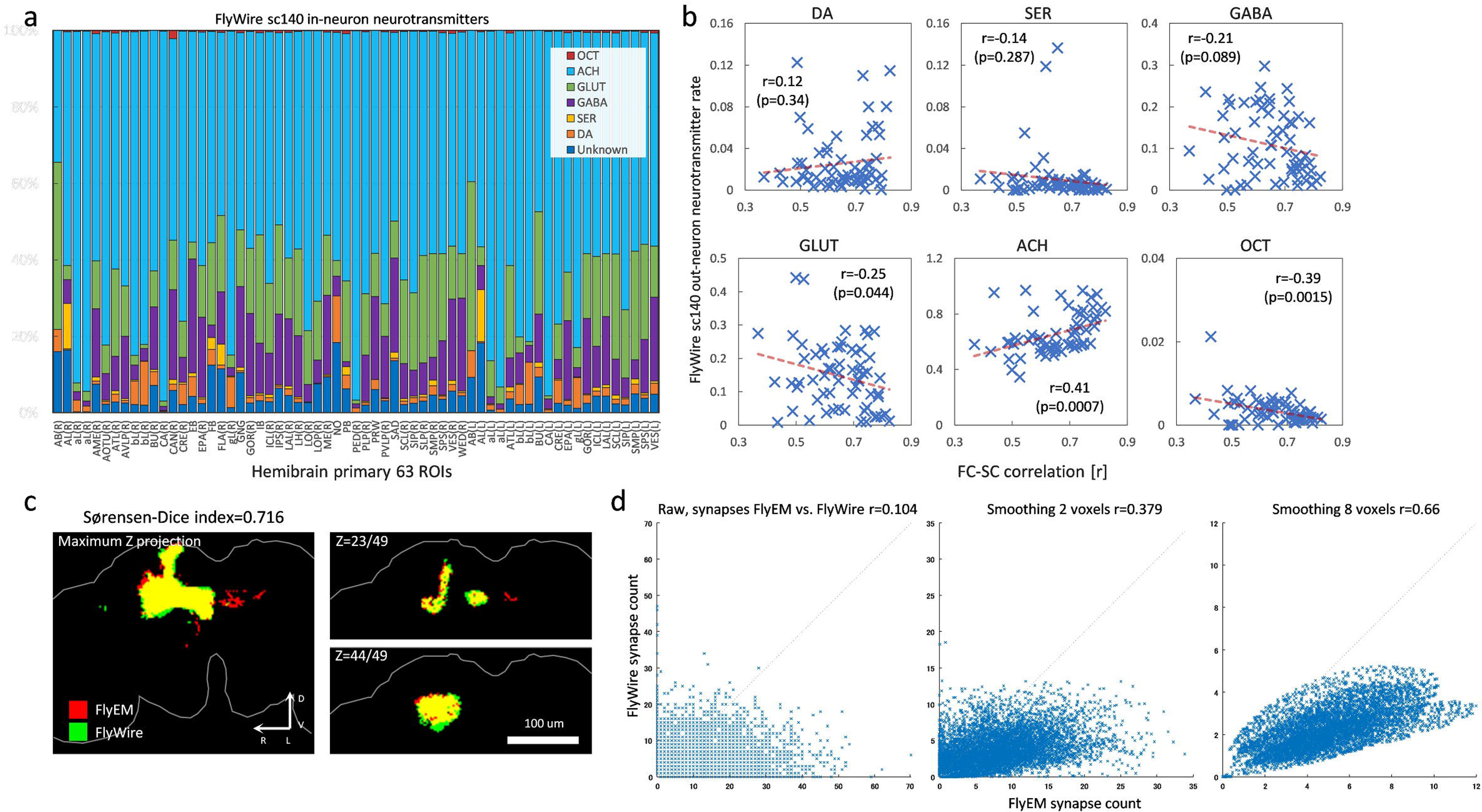
Quantification of pre-and post-synapse segregation and relationship between FC and synapses (FlyEM). a, Unsegregated first (APL_L cell) and second (DPM_R cell) ranked cells in the FlyEM (cf0.8) connectome data. Red dots shows pre-synapses, blue dot shows post-synapses. **b,** The PPSSI histogram of hemibrain neurons (99,644) in FlyEM (cf0.8) connectome data. The vertical axis shows neuron count, and the horizontal axis shows PPSSI [0 1]. **c,** Segregated first (AOTU019_R cell) and second (DNa02_R cell) ranked cells in the FlyEM (cf0.8) connectome data. **d,** Histogram of null SC matrices (blue bar) and extracted SC matrix with cPPSSI (0-0.1) (red solid line). The black dotted line shows the cumulative distribution function of the normal distribution, the brown solid line shows the cumulative distribution function of null & extracted SC matrices, and the red dotted line shows the Bonferroni-corrected p<0.05 threshold. Top shows the SC matrix (synapse) total, middle shows FC-SC correlation (neuron), and bottom shows FC-SC correlation (synapse). **e,** Histogram of null SC matrices and extracted SC matrix with cPPSSI (0.9-1). **f,** Example of a reciprocal-synapse in the EM image (left) and 3D image (right). g, Histogram of reciprocal-synaptic minimum distances in the FlyEM (cf0.8) connectome data. The vertical axis shows reciprocal-synapse pair count, and the horizontal axis shows distance [μm]. **h,** Histogram of null SC matrices and extracted SC matrix with reciprocal-synapses (≤2μm). **i,** The segregation index histogram of all neurons (139,255) in FlyWire (sc140) connectome data. The horizontal axis shows segregation index [0 1]. **j,** The segregation index histogram of hemibrain neurons (99,644) in the FlyEM (cf0.8) connectome data. The horizontal axis shows the segregation index [0 1].

**Extended Data Fig.4-2.**
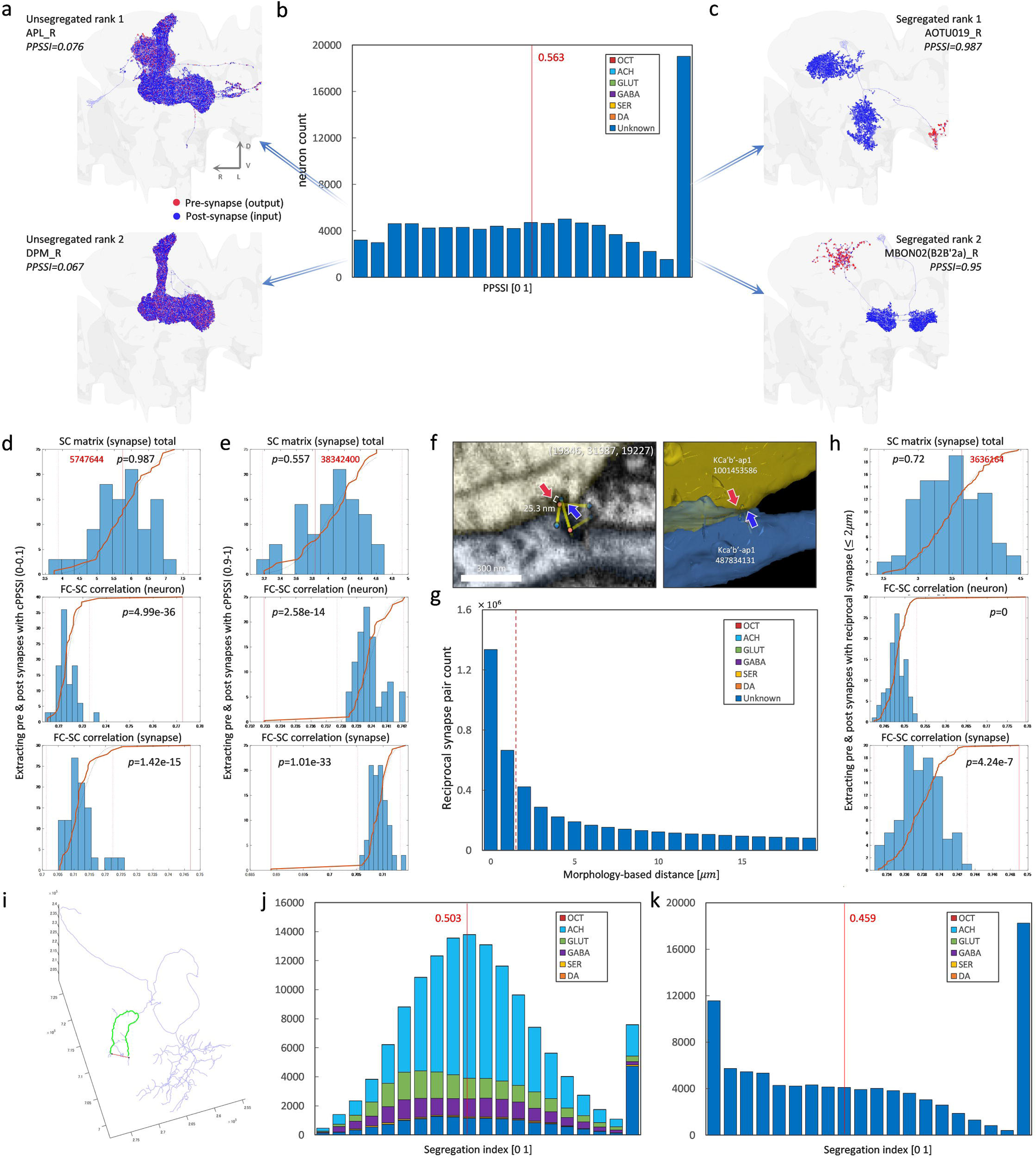
Reciprocal synapses of CT1 and APL neurons, and synapse point cloud for several conditions. a, Histogram of reciprocal-synaptic minimum distances of the CT1-R neuron in the FlyWire (sc140) connectome data. The vertical axis shows reciprocal-synapse pair count, and the horizontal axis shows distance [μm]. **b,** Skeleton view of the CT1-R neuron. Red dots are pre-synapses and blue dots are post-synapses of reciprocal synapses (≤ 2*μm*). **c,** Histogram of reciprocal-synaptic minimum distances of the APL-R neuron in the FlyWire (sc140) connectome data. **d,** Skeleton view of the APL-R neuron, the same way as b. **e,** (top) Maximum Z-projection of pre & post synapse point cloud within cPPSSI (0-0.1) in the FlyEM data (cf0.8). (bottom) Maximum Z-projection of pre & post synapse point cloud within cPPSSI (0-0.1) in the FlyWire data (sc140). **f,** The same as in e, but with the condition cPPSSI in 0.9-1. **g,** The same as in e, but with the condition of only reciprocal synapses (≤ 2*μm*).

**Extended Data Fig.4-3.**
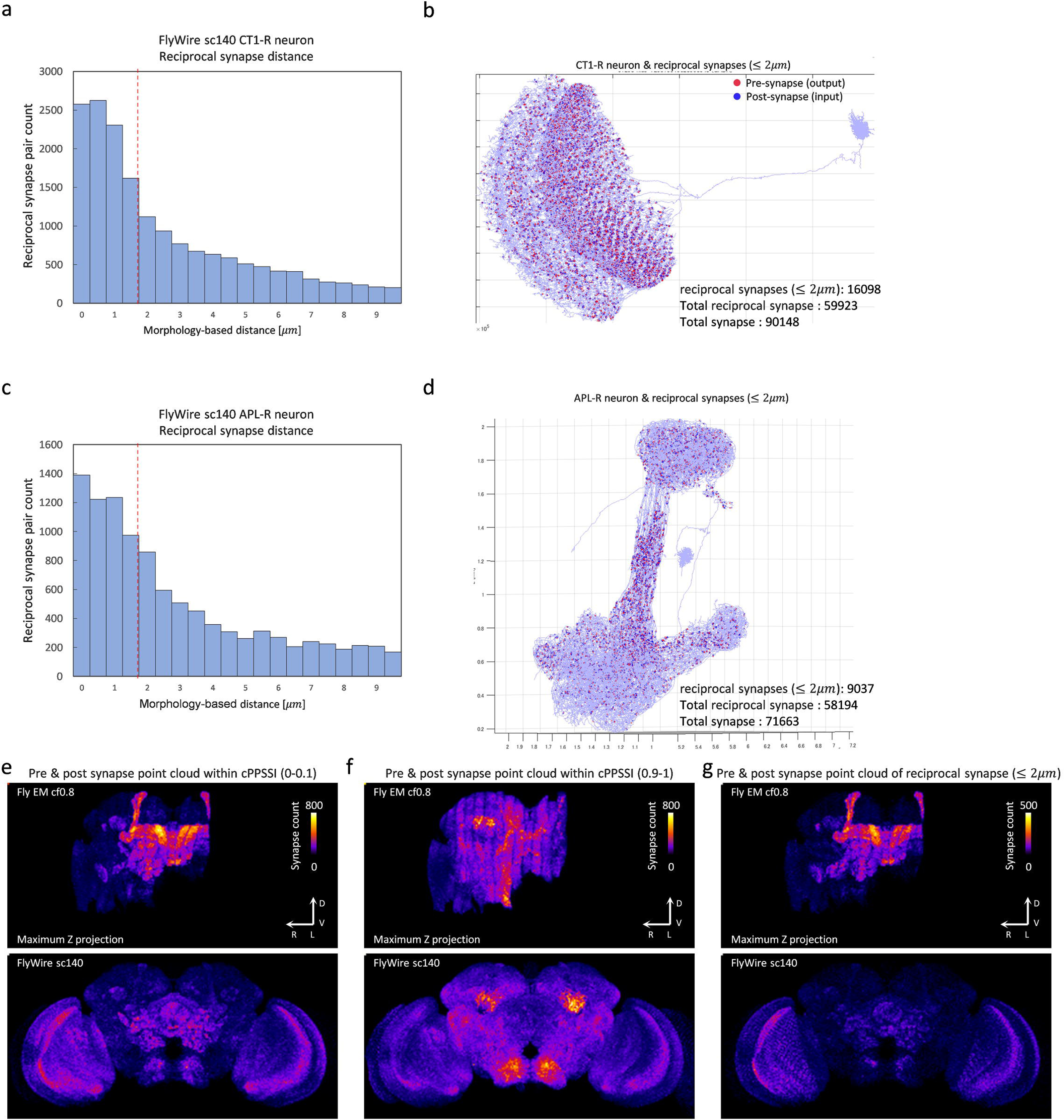
Unsegregated and segregated cell ranking examples (FlyWire sc140). a, Unsegregated cell ranking from 3^rd^ to 11th in the FlyWire (sc140) connectome data. Red dots shows pre-synapses, blue dots shows post-synapses. **b,** Segregated cell ranking from 3^rd^ to 11th.

**Extended Data Fig.4-4.**
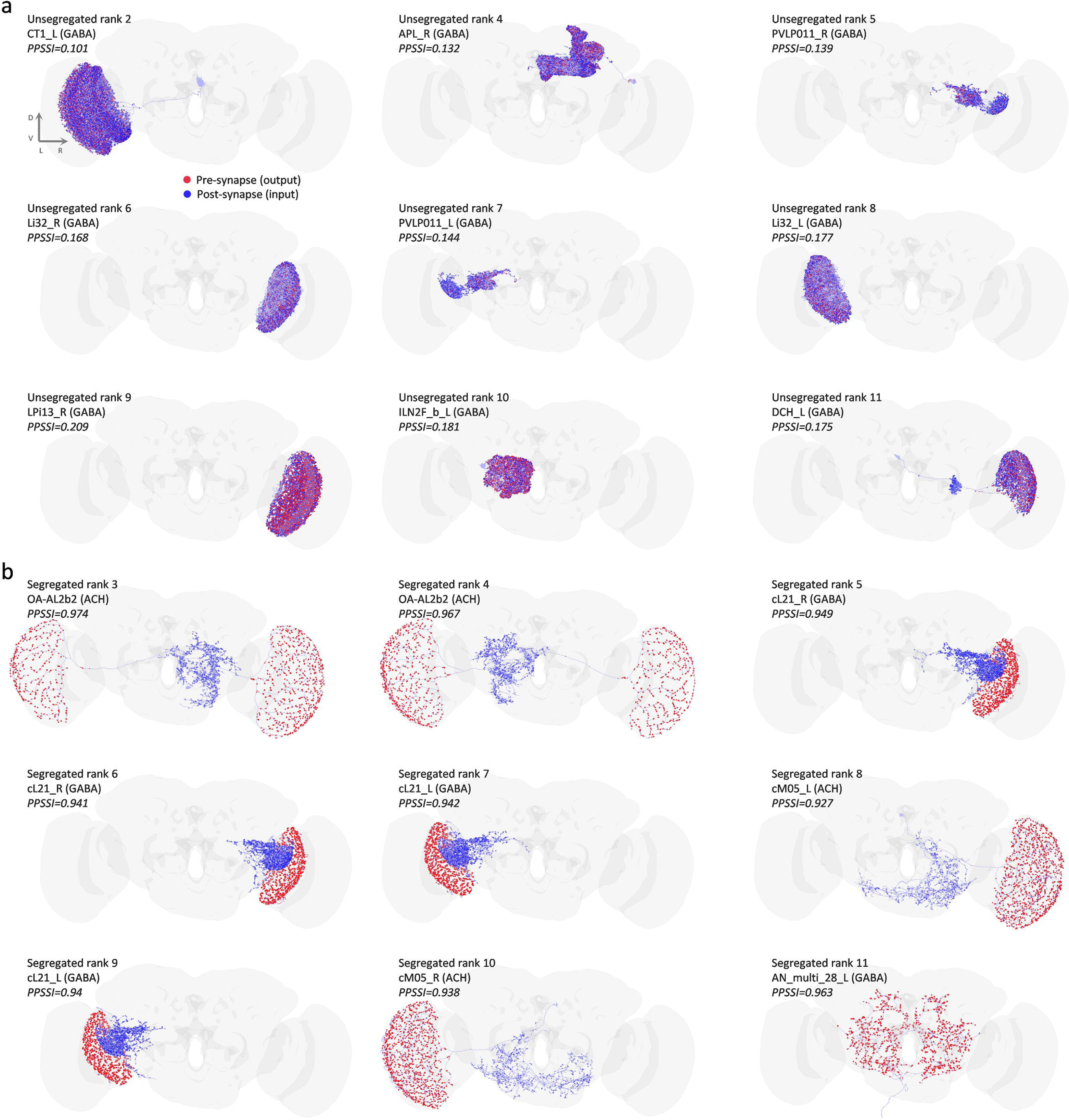
Unsegregated and segregated cell ranking (FlyEM cf0.8). a, Unsegregated cell ranking from 3^rd^ to 11th in the FlyEM (cf0.8) connectome data. Red dots shows pre-synapse, blue dots show post-synapses. **b,** Segregated cell ranking from 3^rd^ to 11th.

**Extended Data Fig.4-5.**
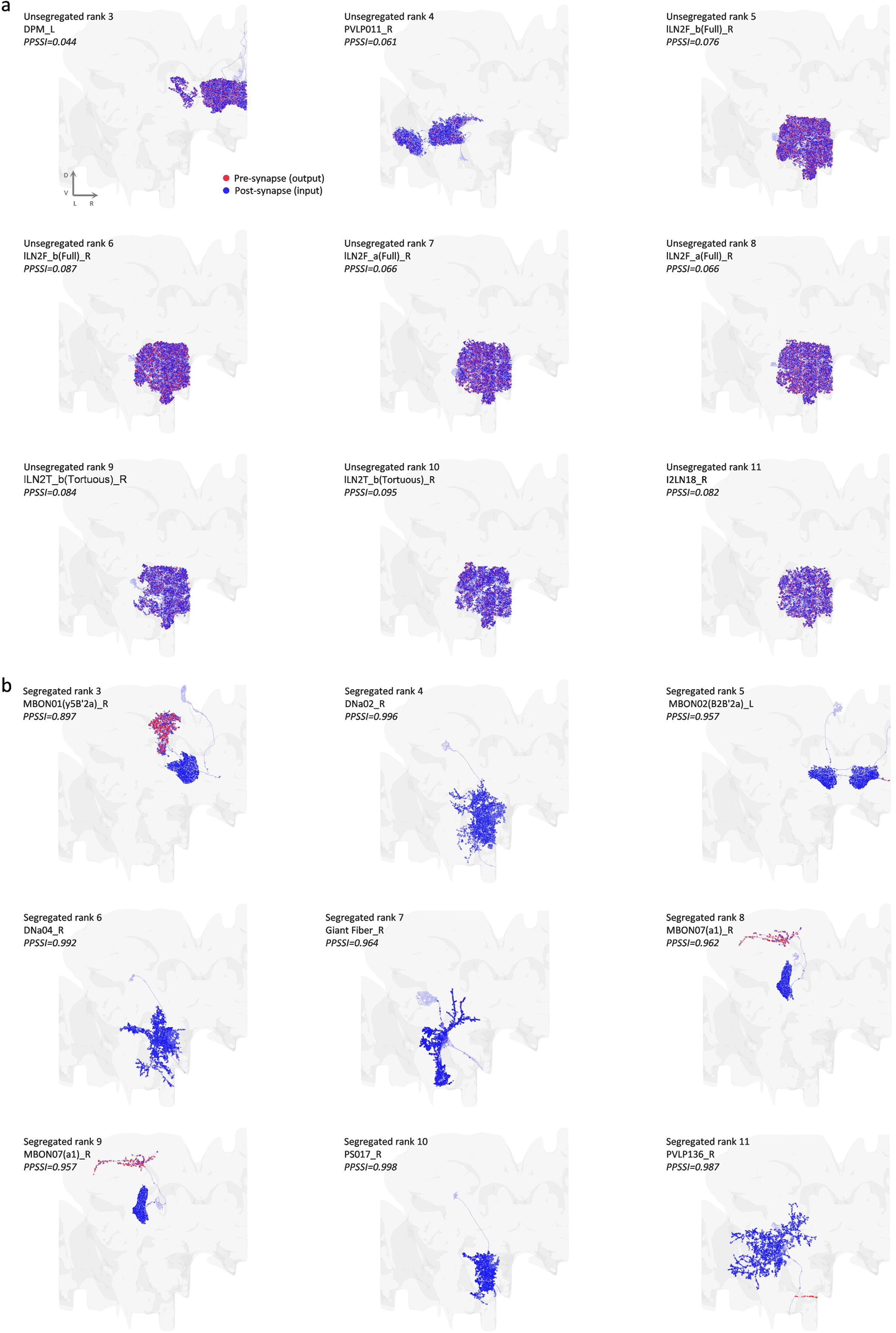
Sparsity rate of extracted SC matrix. **a**, Sparsity rate histogram of SC matrix with cPPSSI (0-0.1) and subsampled null SC matrices corresponding to Fig.4e. The red line indicates sparsity rate of the SC matrix with cPPSSI (0-0.1). **b**, Sparsity rate histogram of SC matrix with cPPSSI (0.9-1) and subsampled null SC matrices, corresponding to Fig.4f. **c**, Sparsity rate histogram of SC matrix with reciprocal synapse (≤2*μm*) and subsampled null SC matrices, corresponding to Fig.4i.

**Extended Data Fig.5-1.**
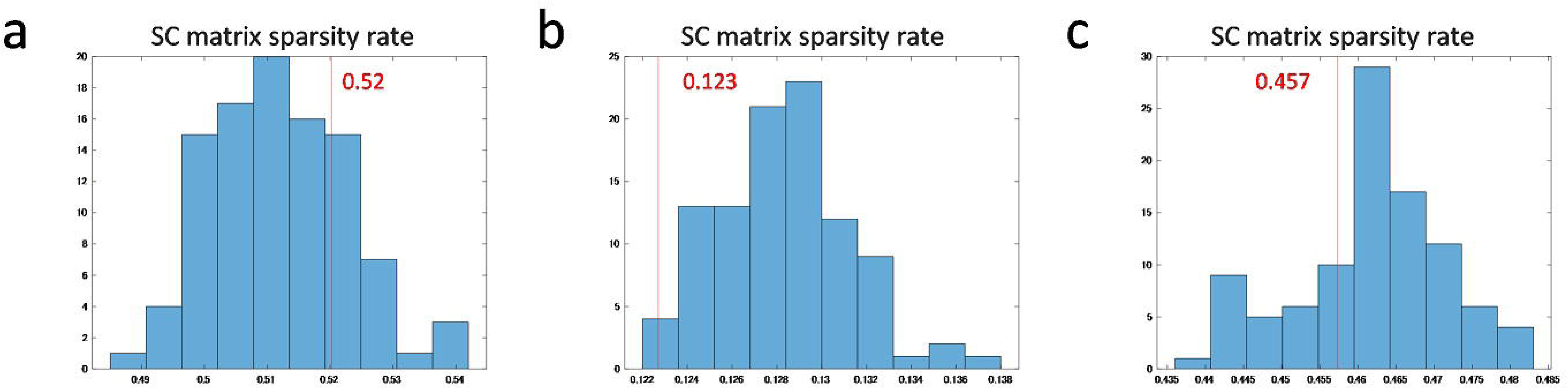
Mechanosensory pathway from Wedge to Ellipsoid Body. a, 3D image of WL-L right, WED016 left, WPNb tier 1, and ER1 left neurons, frontal view. Black square insert shows WL-L right, WED016 left, WPNb tier 1 and ER1 left neurons from the left lateral view. **b,** Network graph of WL-L right, WED016 left, WPNb tier 1 and ER1 left neurons from the FlyWire codex (sc50, connection threshold≥5). Blue is for acetylcholine, ocher for GABA neurons. The small number on the edges shows synapse count.

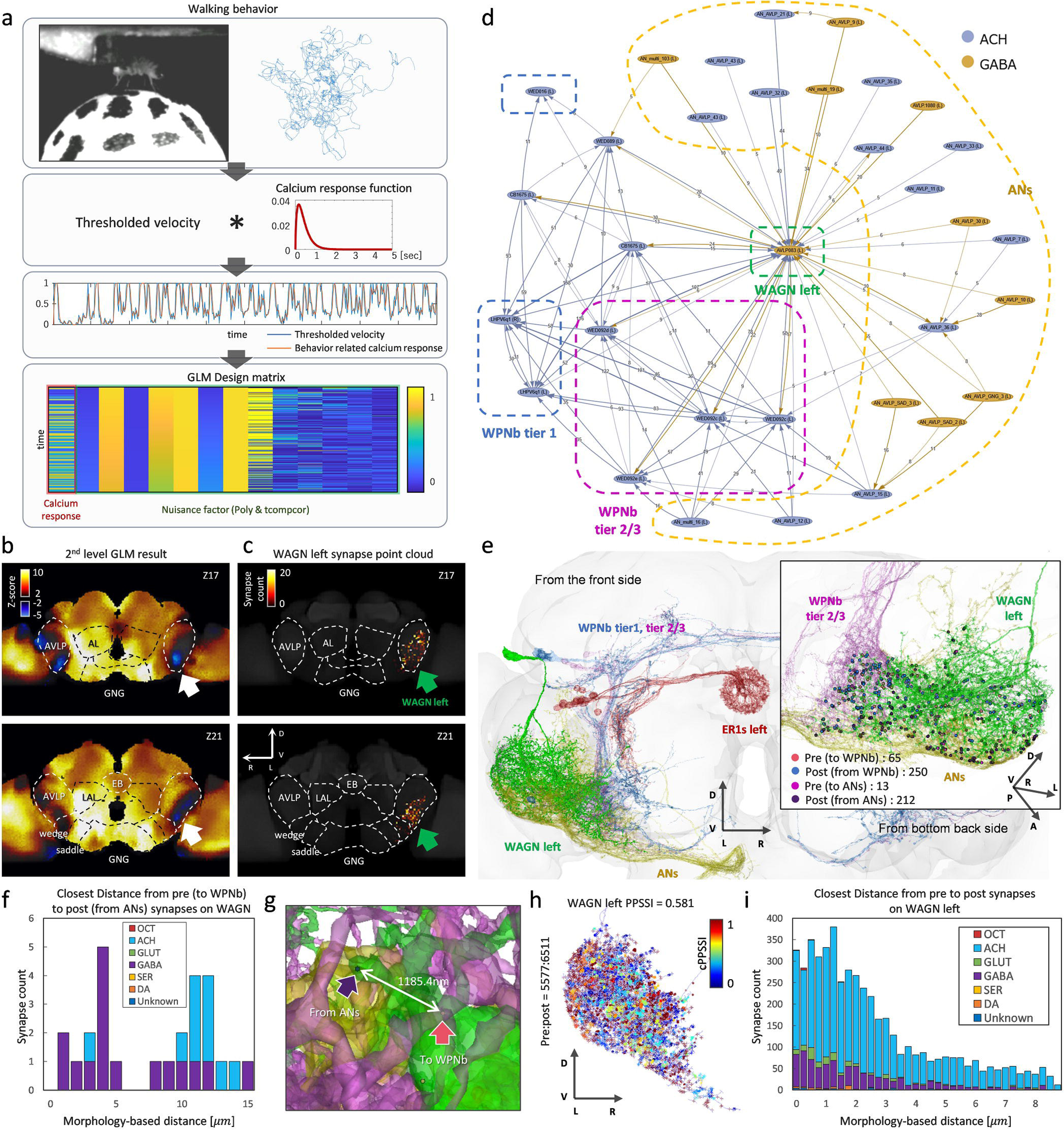

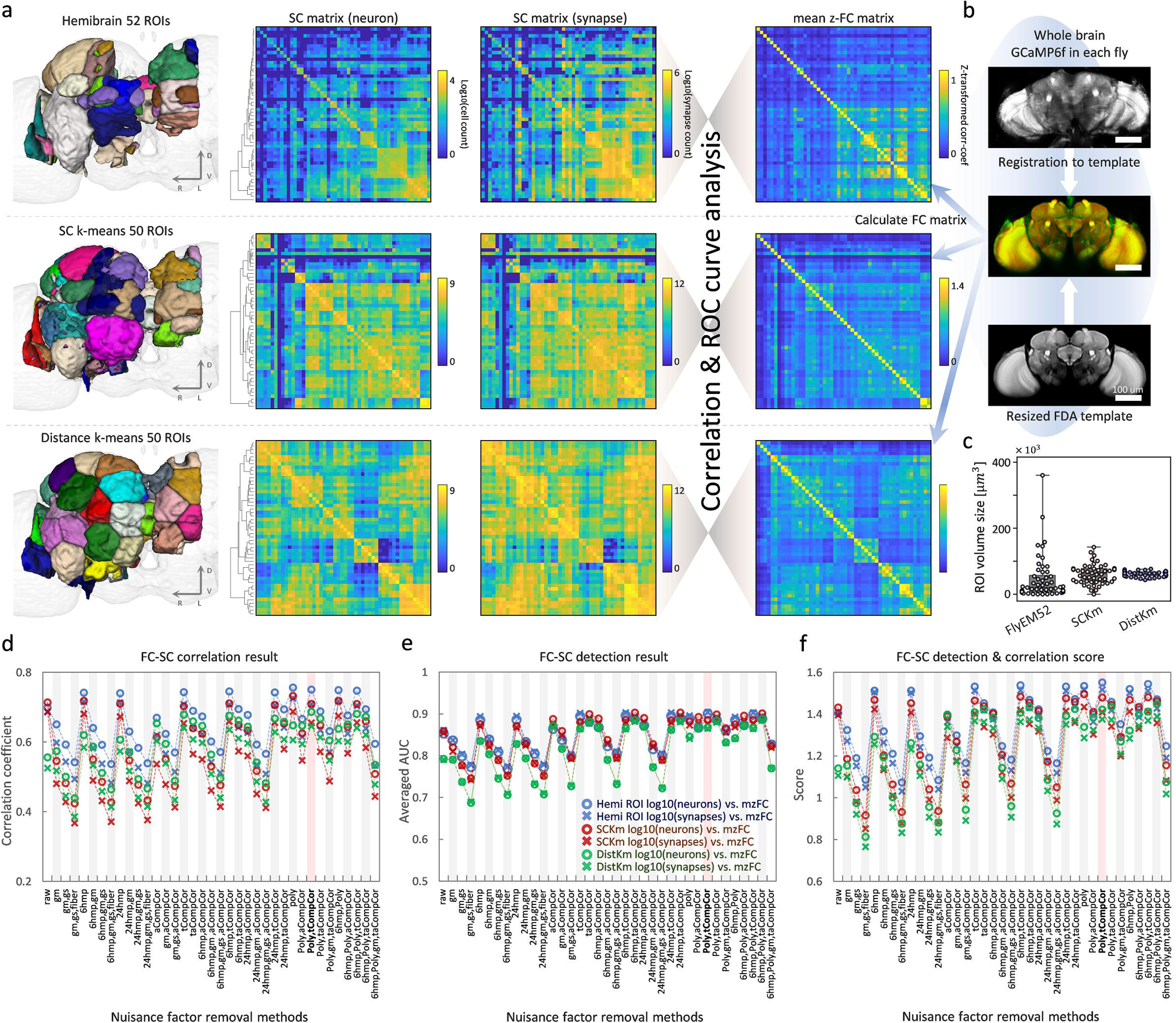

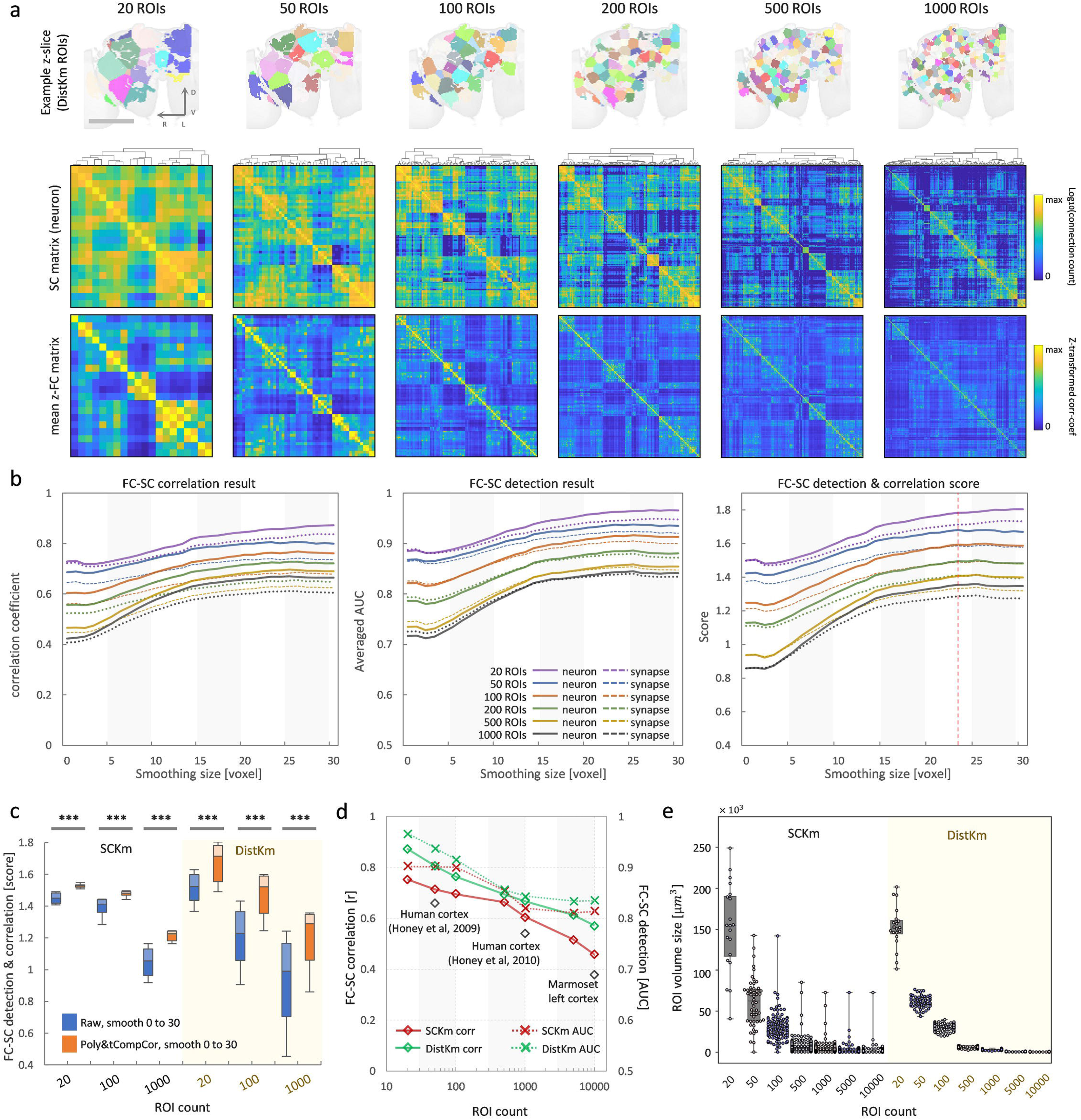

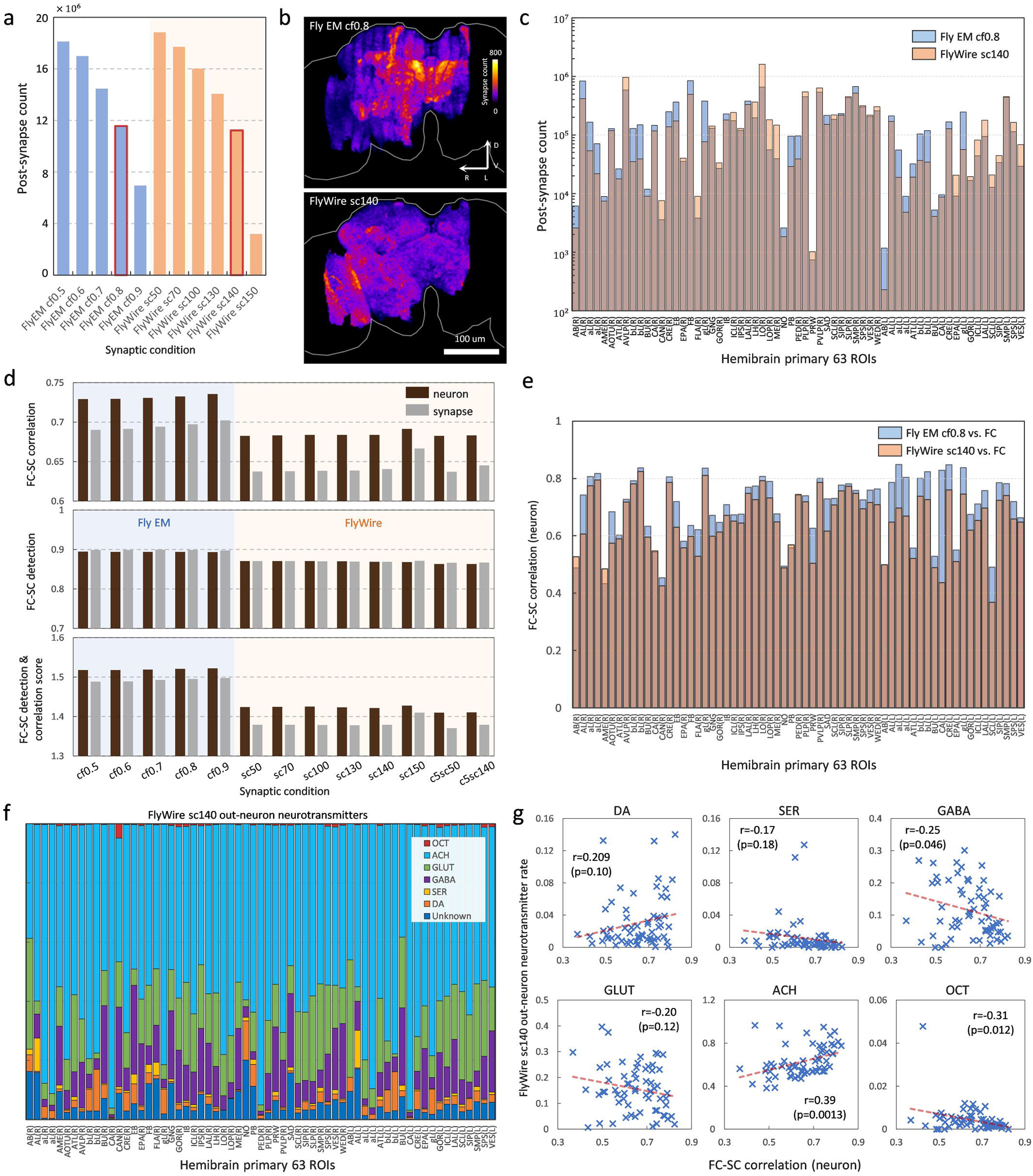

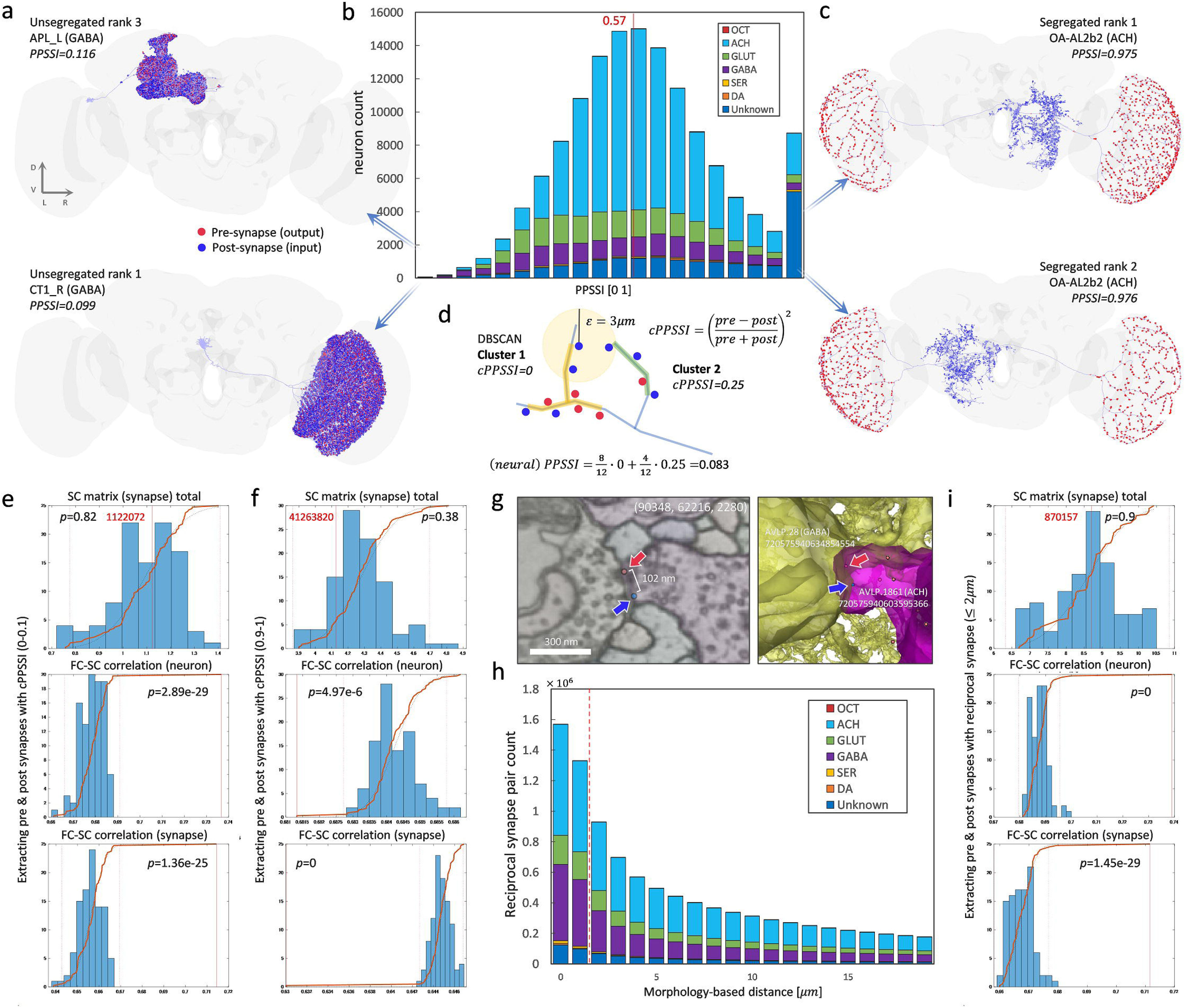

